# The rate of adaptive molecular evolution in wild and domesticated *Saccharomyces cerevisiae* populations

**DOI:** 10.1101/2022.12.07.519429

**Authors:** Maximilian W. D. Raas, Julien Y. Dutheil

**Author notes:** Theoretical Biology and Bioinformatics, Biology, Science Faculty, Utrecht University, 3584 CH Utrecht, The Netherlands.

## Abstract

Through its fermentative capacities, *Saccharomyces cerevisiae* was central in the development of civilization during the Neolithic period, and the yeast remains of importance in industry and biotechnology giving rise to *bona fide* domesticated populations. Here, we conduct a population genomic study of domesticated and wild populations of *S. cerevisiae*. Using coalescent analyses, we report that the effective population size of yeast populations decreased since the divergence with *S. paradoxus*. We fitted models of distribution of fitness effects to infer the rate of adaptive (*ω_a_*) and non-adaptive (*ω_na_*) non-synonymous substitutions in protein-coding genes. We report an overall limited contribution of positive selection to *S. cerevisiae* protein evolution, albeit with higher rates of adaptive evolution in wild compared to domesticated populations. Our analyses revealed the signature of background selection and possibly Hill-Robertson interference, as recombination was found to be negatively correlated with *ω_na_* and positively correlated with *ω_a_*. However, the effect of recombination on *ω_a_* was found to be labile, as it is only apparent after removing the impact of codon usage bias on the synonymous site frequency spectrum and disappears if we control for the correlation with *ω_na_*, suggesting it could be an artefact of the decreasing population size. Furthermore, the rate of adaptive non-synonymous substitutions is significantly correlated with the residue solvent exposure, a relation that cannot be explained by the population’s demography. Together, our results provide a detailed characterization of adaptive mutations in protein-coding genes across *S. cerevisiae* populations.

## Introduction

The domestication of crops and livestock is widely regarded as a defining determinant in the rise of permanent human settlement and the development of contemporary civilisation during the Neolithic period (Diamond, 2002; Fuller & Stevens, 2019; Zeder, 2008). The budding yeast *Saccharomyces cerevisiae* is believed to be among the earliest domesticated species, as archaeological evidence has shown that humans were crafting fermented beverages as early as 9,000 years ago in ancient China (McGovern et al., 2004). From its initial use in liquid fermentation reactions, domesticated usage of *S. cerevisiae* has further diversified into bread leavening and myriad biotechnological applications (Lahue, Madden, Dunn, & Smukowski Heil, 2020; Parapouli, Vasileiadis, Afendra, & Hatziloukas, 2020), and it has been adopted as a leading model organism in molecular biology and (evolutionary) genetics (Botstein & Fink, 2011; Duina, Miller, & Keeney, 2014; Marsit et al., 2017), being the first eukaryote to have its full genome sequenced (Goffeau et al., 1996). The close alliance between humankind and yeast has led to the global dispersal of *S. cerevisiae* through trade and migration, leaving a marked impact on the life history of the yeast (Legras, Merdinoglu, Cornuet, & Karst, 2007; Ludlow et al., 2016; Money, 2018; Sicard & Legras, 2011). Consequently, industrially-used strains of *S. cerevisiae* show distinct patterns of demographic history and genome evolution marked by typical domestication signatures, including historic bottlenecks and higher rates of unbalanced rearrangements, polyploidies and aneuploidies in domesticated compared to wild strains (Duan et al., 2018; Gallone et al., 2016; Liti et al., 2009; Peter et al., 2018; Yue et al., 2017). In contrast to these highly domesticated strains, the relatively recent discovery of truly wild populations of *S. cerevisiae* in Asian primeval forests showed that there are present-day populations that have continued to evolve under purely natural conditions (Liti, 2015; Wang, Liu, Liti, Wang, & Bai, 2012), countering the long-standing view that *S. cerevisiae* is an exclusively domesticated species. High-resolution phylogenetic analyses have confirmed the historic separation between wild and domesticated populations (Duan et al., 2018; Gallone et al., 2016; Legras et al., 2018; Ludlow et al., 2016; Peter et al., 2018). Therefore, while *S. cerevisiae* isolates routinely used in research and industry represent *bona fide* domesticated populations, there also exist truly wild populations of *S. cerevisiae* in their native Far East Asia. Owing to the unique availability of large amounts of genomic data from both these domesticated and wild populations, *S. cerevisiae* provides an outstanding opportunity to investigate the impact of domestication on genome evolution.

The study of domestication has a long-standing tradition in evolutionary biology, and has been influential in establishing the theory of evolution by natural selection (Darwin, 1868). It is known to impact population demographic dynamics substantially and, at the molecular level, genome diversity and evolution. One open question is how domestication impacts the relative contributions of positive selection and genetic drift to genome evolution (Moyers, Morrell, & McKay, 2017). With the dawn of the (post-)genomic era, novel population genomics approaches towards delineating the relative contributions of these drivers of sequence evolution have become available. Importantly, these methods enable the quantification of adaptive sequence evolution (i.e., resulting from positive selection) from non-adaptive sequence evolution (i.e., attributable to genetic drift and purifying selection). This is made possible by the comparison of intra-specific (polymorphism) and inter-specific (divergence) data, an approached pioneered by (Hudson, Kreitman, & Aguadé, 1987) and extended by (McDonald & Kreitman, 1991). The latter approach makes the distinction between two types of mutations in protein-coding sequences: synonymous mutations, which do not change the encoded amino-acid owing to the redundancy of the genetic code and are thus assumed to be neutral, and non-synonymous mutations, which change the amino-acid sequence, possibly altering the function of the resulting protein and making these subject to selection. The comparison of the amount of synonymous and non-synonymous polymorphism on the one hand, with the quantity of synonymous and non-synonymous divergence on the other hand allows estimating the proportion of non-synonymous substitutions fixed by positive selection, a metric termed *α* by (Smith & Eyre-Walker, 2002). More recent developments include the modelling of the distribution of fitness effects (DFE) underlying polymorphism and divergence patterns obtained from population genomics data (Adam Eyre-Walker & Keightley, 2007; A. Eyre-Walker & Keightley, 2009; A. Eyre-Walker, Woolfit, & Phelps, 2006). This approach was later further extended to explicitly account for slightly beneficial mutations (Galtier, 2016; Tataru, Mollion, Glémin, & Bataillon, 2017). These statistical approaches permit the inference of parameters describing historic adaptive evolution: the rate of adaptive non-synonymous substitutions (*ω_a_*), the rate of non-adaptive non-synonymous substitutions (*ω_na_*) and the proportion of adaptive non-synonymous substitutions (*α*).

There are various ways in which domestication could have impacted the contributions of adaptive versus non-adaptive evolution in *S. cerevisiae*. On the one hand, studies of animal and plant domestication have shown that the strong directional selection involved in domestication leaves notable signatures of adaptive evolution on domestication-associated genes (Ahmad et al., 2020; Ross-Ibarra, Morrell, & Gaut, 2007). Therefore, it is possible that similar mechanisms underly niche-adaptation in domesticated yeast populations (Duan et al., 2018; Legras et al., 2018), in turn yielding elevated values for *ω_a_* in genes involved in phenotypic adaptations towards domesticated niches. On the other hand, differences in adaptive evolution across populations may be the result of their distinct effective sizes. The effective population size (*N_e_*) and the rate of adaptive evolution are tightly linked as, in small populations, (i) genetic drift is stronger, preventing advantageous mutations with small effect (lower than 1/Ne) to get fixed, and (ii) the available supply of standing genetic variation will generally be lower (Charlesworth, 2009; Gossmann, Keightley, & Eyre Walker, 2012; Lanfear, Kokko, & Eyre-Walker, 2014). Importantly, domestication is associated with strong demographic dynamics and domestication-associated bottlenecks have been identified in domesticated *S. cerevisiae* populations (Gallone et al., 2016). Therefore, it is expected that domesticated populations of *S. cerevisiae* have lower overall rates of adaptive evolution compared to wild populations as a result of the domestication bottleneck, despite the strong directional selection acting on specific cellular processes.

Another major driver of the rate of evolution (both adaptive and non-adaptive) is recombination, which impacts the efficacy of both adaptive and purifying selection as it can combine interfering adaptive mutations and unlink adaptive mutations from deleterious ones (Marais & Charlesworth, 2003). The relation between sexual reproduction and the efficacy of selection is well-documented in yeasts (Connallon & Knowles, 2007; Goddard, Godfray, & Burt, 2005). It has recently been reported that domesticated populations of *S. cerevisiae* largely abolish sexual reproduction (De Chiara et al., 2022) and do not undergo meiotic recombination. Therefore, the suppression of recombination in domesticated populations is also expected to induce a lower rate of adaptive evolution in domesticated populations.

Earlier landmark work by (Liti et al., 2009) identified no evidence for adaptive evolution in a set of ∼1,100 *S. cerevisiae* genes from a diverse set of *S. cerevisiae* strains, using the method developed by (McDonald & Kreitman, 1991). However, this method has important limitations when it comes to accounting for slightly deleterious mutations, which can therefore substantially bias inferences of adaptive evolution (A. Eyre-Walker & Keightley, 2009). Moreover, protein structure plays an important role in shaping the rate of substitutions. Specifically, residues with a high relative solvent accessibility (RSA) show higher rates of both adaptive and non-adaptive substitutions (Moutinho, Trancoso, & Dutheil, 2019). Accounting for RSA will yield higher resolution insights into the factors shaping substitution rates in *S. cerevisiae* and allow for more sensitive detection of adaptive sites.

Here, we use a population genomics approach to assess the impact of several factors on the rate of adaptive and non-adaptive non-synonymous substitutions in wild and domesticated populations of *S. cerevisiae*. By comparing 17 populations of different origins, we show that domestication resulted in both a decrease of the rate of adaptive – and an increase of the rate of non-adaptive – non-synonymous substitutions, and that the effective population size is a determinant of both rates. We find that only the rate of non-adaptive non-synonymous substitutions is significantly lower in the domestication-associated functional categories of genes. Adaptive non-synonymous substitutions are more frequently found outside the core proteome, that is, in isolate-specific genes. Signatures of linked selection are pervasive, with recombination rate being negatively correlated with *ω_na_* and positively correlated with *ω_a_*. This effect, however, is weak and masked by codon usage bias when not accounted for and disappear if we correct for the correlation of recombination with the strength of purifying selection. Finally, the residues’ solvent exposure strongly impacts both adaptive and non-adaptive rates, with the majority of adaptive substitutions occurring at the protein surface.

## Materials & Methods

### Genomic dataset and population definition

The *S. cerevisiae* genome data used in this study was generated by (Peter et al., 2018) as part of the “1,002 Yeast Genomes Project”. Unless specified otherwise, the *de novo* genome assembly of each isolate was used in the following analyses. Relevant populations of isolates were defined based on the clades as determined in the original publication, followed by manual filtering of isolates based on ecological and geographical origins to assure population homogeneity. We included populations with a minimum sample size of five isolates for further analyses. The Italian Wine population had a large sample size and it was randomly split into two subsets of isolates for computational tractability. Given that results were highly similar between the two subsets, we only report the ones for one subset, which we refer to as the “Italian Wine” population in the following. Combined, this yields a total of 17 *S. cerevisiae* populations from both domesticated and wild origins (*Supplementary Table 1*).

### Genome alignment and gene extraction

For each population, we generated full genome alignments and extracted the protein-coding DNA sequence (CDS). The genomes in each population and the reference genome R64 (Downloaded from ftp://ftp.ensemblgenomes.org/pub/fungi/release-49/fasta/saccharomyces_cerevisiae/dna/; last accessed December 2020) were aligned using the MultiZ v11.2 alignment program from the Threaded-Blockset Aligner tool suite (Blanchette et al., 2004). The alignments were then projected onto the reference sequence, and the resulting alignment blocks were realigned and filtered using MafFilter v1.3.1 (Dutheil, Gaillard, & Stukenbrock, 2014). Alignment blocks were split into windows with a maximum size of 10 kb, which were subsequently realigned using MAFFT v7.38 (Katoh & Standley, 2013). We then filtered the alignment to only include i) blocks with sequences of all selected isolates in the population and ii) blocks with a minimum length of 10 bp. Ten bp windows, slid by 1 bp, containing ≥ 30% gap and unresolved characters were discarded. The remaining alignment blocks were merged if not further apart than 100 bp and between-block positions were filled with unresolved characters “N”, resulting in a masking of ambiguously aligned regions. A callability mask was also generated, recording the coordinates of the filtered positions. Diversity statistics (average pairwise heterozygosity) were computed in 10 kb non overlapping windows along the alignment, for each population.

CDSs were extracted according to the General Feature Format (GFF) file of the reference sequence (Downloaded from ftp://ftp.ensemblgenomes.org/pub/fungi/release-49/gff3/saccharomyces_cerevisiae; last accessed December 2020) and the obtained CDSs were concatenated to reconstruct genes from the alignment files. Reconstructed gene-coding sequences were checked for premature stop codons to ensure accurate reconstruction. No genes were removed at this step. In these and subsequent procedures, genomic data was handled with scripts using the Biopython package v1.78 (Cock et al., 2009), available at https://gitlab.gwdg.de/molsysevol/saccharomycespopgen. The total number of nucleotide sites and final number of genes for each population are summarised in *Supplementary Table 2*.

### Phylogenetic tree

We used the complete (unaligned) genome sequences of all isolates in the selected populations to compute pairwise distances using *andi* v0.12 (Haubold, Klötzl, & Pfaffelhuber, 2015). A phylogenetic tree was inferred from the resulting distance matrix using FastME v2.0 (Lefort, Desper, & Gascuel, 2015) and plotted using FigTree v1.4.4 (Rambaut, 2018).

### Demographic history inference

We used the Multiple Sequentially Markovian Coalescent 2 v2.1.1 (MSMC2) (Malaspinas et al., 2016) to reconstruct the demographic histories of the selected populations. As SMC-based methods have been benchmarked on Primates dataset (Schiffels & Wang, 2020), we conducted a simulation analysis to assess their accuracy on yeast datasets (see *Supplementary Methods*). We assessed the power of the MSMC2 method by simulating data under a neutral model of sequence evolution, which fulfilled the hypotheses of the inference model (homogeneous recombination and mutation rate, no selection). We simulated 10 datasets of 5 diploid individuals (10 haploid genomes), with the same chromosome structure as the reference *S.cerevisiae* genome (R64). We found that the default parametrisation of the time epochs led to biased estimates in the most recent and ancient times (*Supplementary Figure 1A*). We then fitted a model with fewer parameters. We considered 30 epochs, with the five most recent and the five most ancient epochs sharing a common coalescence rate, leading to 22 parameters, the 20 intermediate epochs being allowed to have a distinct coalescence rate. With this discretisation scheme, no bias was observed (*Supplementary Figure 1B*). However, we observed an increased estimation variance in the most recent and ancient times. Adding a distribution of missing data identical to the one of one of our population alignments (Far East Asia) did not significantly affect the inference (*Supplementary Figure 1C*, repeated in *Figure 3A*). For all remaining MSMC2 inferences, we used the 22 parameters time pattern.

We separated the dataset into isolates for which the *de novo* assembled genome could be considered as a haplotype (haploid and diploid homozygous isolates) and isolates for which the assembled genome represented a consensus genome (diploid heterozygous isolates). In the first case, MSMC-compatible input files were generated from the filtered genome alignments using MafFilter v1.3.1 (Dutheil et al., 2014). In the second case, the corresponding diploid genotypes were extracted from the 1,011 yeast genome variant calls using the bcftools (Danecek et al., 2021; Peter et al., 2018). Positions with missing genotypes were masked. As a callable mask, we used the positions of the *de novo* genome alignments, after filtering. Because the variants are unphased, only haploid pairs from the same isolate were compared in MSMC2 (n pairs, where n is the number of isolates in a population), while all haplotype pairs were analysed in the case of haploid or homozygous diploid isolates (n*(n-1)/2 pairs). Following our benchmark procedure, MSMC2 was run using a 1*5+20*1+1*5 discretisation scheme. Other options were kept as program defaults. We ran MSMC2 on all isolates in each population, but also separately on several pairs of haplotypes to estimate the robustness of the inferred demography for each population, as well as the homogeneity of the samples. Results are reported in coalescent units (x-axis) and inverse instantaneous coalescence rate (IICR, y-axis).

We estimated the effective population size of the *S. cerevisiae* / *S. paradoxus* ancestor using the IM-CoalHMM model (Mailund et al., 2012). *S. paradoxus* is a close relative of *S. cerevisiae* and was for this reason selected as the outgroup for this analysis. The model considers a speciation process with an initial differentiation phase with symmetric migration between the two populations, followed by a phase where the two populations are genetically isolated. The estimated parameters are the start and end of the migration phase, the average ancestral population size, and the average recombination rate. A seven species genome alignment of yeasts was downloaded from the UCSC genome browser. The alignment was projected on the *S. cerevisiae* (sacCer3) genome using the ‘maf_project’ program v12 from the TBA package (Blanchette et al., 2004) and the pairwise alignment of *S. cerevisiae* with *S. paradoxus* (sacPar) was extracted and merged, replacing non-aligned positions by missing data. The 16 chromosomes were further concatenated before being input to the IMCoalHMM program (v0.7.0, https://github.com/mailund/IMCoalHMM).

### Quantifying adaptive evolution

For reconstructed gene alignments, an outgroup sequence was identified using the closest match of a ‘blastn’ v2.9.0+ search (Altschul, Gish, Miller, Myers, & Lipman, 1990) on the *S. paradoxus* genome (downloaded from https://ftp.ncbi.nlm.nih.gov/genomes/refseq/fungi/Saccharomyces_paradoxus/latest_assembly_versions/GCF_002079055.1_ASM207905v1/). Specifically, the reference sequence of each *S. cerevisiae* gene alignment (R64) was queried against the reference CDS file of *S. paradoxus* (Downloaded from https://yjx1217.github.io/Yeast_PacBio_2016/data/) (Yue et al., 2017). Protein-coding sequences, including the outgroup sequence, were re-aligned using the codon alignment program MACSE v2 (Ranwez, Douzery, Cambon, Chantret, & Delsuc, 2018). We calculated statistics on synonymous and nonsynonymous divergence between the species, transversion per transition ratio, and inferences on ancestral alleles from these alignments using the BppPopStats v2.4.0 program from the Bio++ Program Suite (Guéguen et al., 2013) and used them to produce unfolded site frequency spectra (SFSs). Models of distribution of fitness effects (DFE) were then fitted with Grapes v1.1 (Galtier, 2016), to estimate the rates of adaptive and non adaptive non-synonymous substitutions (*ω_a_* and *ω_na_*), and the proportion of adaptive non-synonymous substitutions (α). Grapes was run with the ‘-no_div_param’ option, as the underlying model here was shown to be more robust in light of historic changes in population size (M. Rousselle, Mollion, Nabholz, Bataillon, & Galtier, 2018). Additionally, the option ‘-nb_rand_start 5’ was specified to test distinct optimization starting conditions and prevent model settling on suboptimal local maximum likelihoods. Three DFE models were fitted onto the data: GammaExponential (GammaExpo), DisplacedGamma (DisplGamma) and ScaledBeta (Galtier, 2016). To determine the quality of model fit, Akaike’s Information Criterion (AIC) was calculated for the complete SFSs per population (Akaike, 1973). Here, the GammaExpo and ScaledBeta models performed comparably well, but the ScaledBeta model failed to estimate *ω_a_* in more cases than the GammaExpo model. Therefore, the GammaExpo model was determined as the best performing model, which is in accordance with previous results obtained from data in animals (Galtier, 2016), plants (Moutinho et al., 2019) and fungi (Grandaubert, Dutheil, & Stukenbrock, 2019). Finally, sites were bootstrapped to create 150 SFS replicates for each dataset enabling a more comprehensive analysis of α, *ω_a_* and *ω_na_*. For some replicates, optimisation failed and the corresponding replicates were discarded. The minimum number of replicates for which estimates could be obtained was 90 for the North American Oak population in the Metal Stress gene category. The average number of successful bootstraps across all runs is 149.2. The mean values of the estimates over all bootstrap replicates were used as the final estimates of these parameters in each analysis.

### Gene set definition

Gene sets were defined based on “Genes and gene products” annotations obtained from AmiGO 2 (http://amigo.geneontology.org/amigo) using the search terms “Meiosis” and “Sporulation” (corresponding to the gene set “Meiosis”), “Osmotic Stress”, “Oxidative Stress”, “Glucose OR Xylose OR Maltose OR Fructose OR Sucrose” (corresponding to the gene set “Domestic Carbon”), and “Heat Stress” and “Temperature” (corresponding to the gene set “Heat Stress”). The “Metal Stress” gene set was compiled based on data from genetic screens performed by (Thorsen et al., 2009) for arsenic exposure and by (Jo et al., 2007) for copper exposure. Genes that were found in none or multiple of the aforementioned categories were combined into the “Other Genes” and “Multiple sets” categories. All gene sets are restricted to only include the genes that were recovered from all populations, which totalled 3,215 genes (*Supplementary Figure 2*). Gene overlap analysis was performed with the ‘UpSetR’ package v.1.4.0 (Conway, Lex, & Gehlenborg, 2017).

### Recombination landscape

Recombination rates across the genome for *S. cerevisiae* were taken from (Liu, Maclean, & Zhang, 2018), who quantified crossover (CO) rates in a window-based approach with a window size of 20 kb, and these rates were mapped onto the genes included in our analyses based on the genomic location of the midpoint of these genes in the respective reference genomes. The genes were then binned into 15 approximately equal sized categories, except for the populations Ecuadorian, Malaysian and North American Oak, for which the overall genetic variation was lower and so they were binned into 5 categories with sufficient polymorphic sites.

### Prediction of relative solvent accessibility

The relative solvent accessibility (RSA) per site was predicted using NetSurfP-2.0 (Klausen et al., 2019). NetSurfP-2.0 was run on R64 translated CDSs to obtain the RSA for each site. These per site RSA values as computed for R64 were mapped onto the corresponding sites of the population specific alignments. The sites were binned into 15 equal-sized bins based on RSA except for the populations Ecuadorian and Malaysian, which were binned into 5 categories, and the North American Oak population, which was binned into 3 categories.

### Linear modelling

All statistical analyses were conducted in the R software, version 4.1.0 (R Development Core Team, 2021). We fitted generalised least squares (GLS) linear models to assess the impact of several factors on the rate of adaptive and non-adaptive non-synonymous substitutions while accounting for the phylogenetic relationships between populations. Models were fitted with the maximum likelihood method to compute values of Akaike’s information criterion, using the ‘gls’ method of the ‘nlme’ package (Pinheiro, Bates, DebRoy, Sarkar, & R Core Team, 2021) with phylogenetic correlation structures taken from the ‘ape’ package (Paradis & Schliep, 2019). We fitted separate models to each rate (response variable) including estimates for all populations. All models included the life history of the population (Wild, Domesticated-Asian, and Domesticated Non Asian), the culture method (Environment, Repitching, and Starter-based), and the mean synonymous diversity (*π_N_*) as explanatory variables. We specified a correlation model using the previously estimated population phylogenetic tree and fitting branch lengths using the method of Grafen (Grafen, 1989). Residues were normalised using the Box-Cox transform, modified to allow for non-strictly positive values, as implemented in the ‘car’ package (Fox & Weisberg, 2018). To assess the effect of gene sets, recombination rates and RSA, we fitted extended models including each of these variables in addition to life history, culture method, and *π_N_*, together with their interaction. For instance, the model assessing the impact of RSA on *ω_a_* is described by the formula:

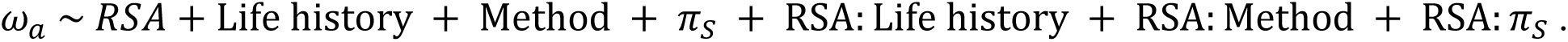

After normalisation of the residues, a model selection was conducted using Akaike’s information criterion (AIC), as implemented in the ‘stepAIC’ function of the ‘MASS’ package (Venables, 2002). Normality and independence of the residues were assessed using the Shapiro and Ljung Box test, as implemented in the ‘stats’ package.

In some cases, the non-normality of the residues was still significant after using the Box-Cox transform, owing to the distribution of *ω_a_*being zero-inflated. Therefore, for comparison, we fitted another set of models where all points with *ω_a_* < 1*e* − 3 were discarded. While these models were fitted on reduced datasets, their residues were generally in better agreement with the Gauss-Markov assumption and had possibly more accurate P-values. Results were visualised with the ‘ggplot2’ (Wickham, 2016), ‘cowplot’ (Claus O. Wilke, 2020), and ‘ggpubr’ (Alboukadel Kassambara, 2020) packages.

### Codon usage analyses

We computed codon frequencies for the set of genes used to estimate rates of adaptive and non adaptive substitutions, for all populations. All 61 non-stop codons frequencies were computed for each analysed site in each population. We did not observe any significant difference in codon usage between populations and summed all counts of all populations. Frequencies were then further summed according to site groups, one of gene functional category, recombination rate category, or RSA category. The resulting contingency tables were submitted to a within-group correspondence analysis as described in (D. Charif, Thioulouse, Lobry, & Perrière, 2004). Relative synonymous codon usage (RCSU) values were also computed for each codon. The RSCU value of a codon is defined as the ratio of the observed codon frequency to the expected frequency if all synonymous codons for the encoded amino-acid were equally used. It is a measure of codon usage bias, which takes values above 1 is the codon is preferentially used, and below 1 if the codon is less preferentially used. The mean codon usage bias for a category of sites was thus computed as the average of the absolute values of RSCU – 1. All codon analyses were conducted using the ‘seqinr’ (Delphine Charif & Lobry, 2007) and ‘ade4’ (Thioulouse, 2018) packages for R.

## Results

We defined 17 population encompassing 329 *S. cerevisiae* isolates from the *1,002 Yeast Genomes Project* genome resource (Peter et al., 2018), based on geography and ecology (*Supplementary Table 1*). A phylogenetic analysis of the selected isolates confirmed that the defined populations represent distinct lineages (*Figure 1*). In agreement with previous reports (Peter et al., 2018), we observed that the domesticated populations formed two distinct groups, based on their Asian or Non-Asian origin (see *Supplementary Results* for a detailed description of the obtained phylogeny). For each population, we estimated the rate of adaptive non-synonymous substitutions in protein-coding genes. We gained insights into the drivers of adaptive evolution between populations and along the yeast genome by estimating rates of adaptive evolution in gene sets with distinct functional categories or local recombination rates, as well as within sites with different solvent exposure. We then compared these estimates between populations, accounting for their evolutionary relationship and demographic history.

**Figure 1:**
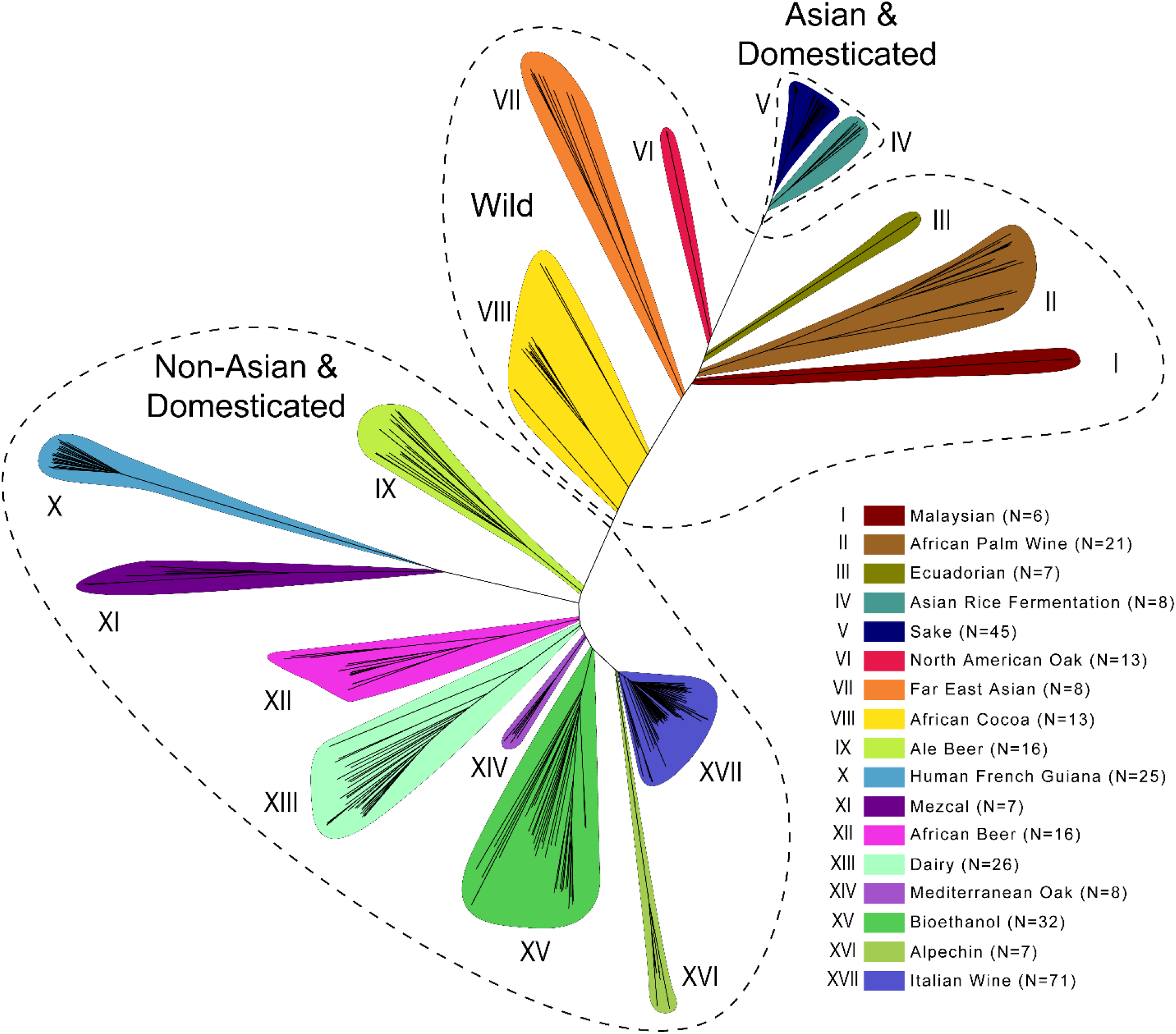
Phylogenetic tree of S. cerevisiae populations. Unrooted phylogenetic tree highlighting the distinct lineages considered as populations in this study. Life history clusters are indicated with dotted lines. Colours indicate population identity as specified in the in-figure legend and numerals.

### A low rate of adaptive non-synonymous substitutions in domesticated *S. cerevisiae*

To study the rate of adaptive evolution in both domesticated and wild *S. cerevisiae* populations, we calculated unfolded site frequency spectra (SFSs) capturing within-population segregating frequencies of synonymous and non-synonymous polymorphic sites. We then fitted models of distributions of fitness effects (DFEs) onto the SFSs using the *Grapes* method (Galtier, 2016), allowing for the computation of the proportion of adaptive non-synonymous substitutions (α), the rate of adaptive non-synonymous substitutions (*ω_a_*), and the rate of non-adaptive non-synonymous substitutions (*ω_na_*). To ensure that the same genes are compared between groups, we restricted our analyses to genes that we could successfully recover from genome alignments in all populations, totaling 3,215 core genes (see *Supplementary Figure 2* and Materials & Methods).

We classified the 17 populations according to two criteria, which we refer to as “life history” and “method”. The first distinguishes wild populations from domesticated populations and within those, the Asian domesticated from non-Asian domesticated populations, as the latter two are believed to be the result of independent domestication events (Duan et al., 2018; Peter et al., 2018). The second criterion refers to distinct application-specific practices for introducing yeasts to fermentation reactions in domesticated settings. These can be broadly classified into “repitching” (also known as “backslopping”) and “starter-based” approaches. In repitching, yeasts are seeded into multiple consecutive fermentation cycles, either by purposely isolating yeasts from an ongoing or terminated fermentation reaction and re-seeding these into fresh wort (traditional repitching, characteristic for beer brewing) (Large et al., 2020), or indirectly via contaminated surfaces that carry residual yeast from previous fermentations such as flasks, baskets or tools (De León- Rodríguez, González-Hernández, Barba de la Rosa, Escalante-Minakata, & López, 2006; Djeni et al., 2020; Schwan & Wheals, 2004). In contrast, yeast cells used in starter-based methods are discarded after a single fermentation reaction. Previous studies have shown that repitching practices significantly affects rates of chromosomal rearrangement rates in *S. cerevisiae*, revealing the substantial impact of application-specific practices on yeast genome evolution (Large et al., 2020). Consequently, we hypothesise that these different practices might impact the rate of adaptation, with repitching methods being likely to better facilitate adaptation through the extended multi-generational exposure to environmental pressures, compared to the continuous evolutionary “reset” of populations resulting from the use of fixed starter cultures.

These analyses revealed that the proportion of adaptive substitution, α, is generally low in *S. cerevisiae* (*Table 1*), with some differences between wild and domesticated populations. Focusing on core genes present in all populations, α values range from 0 (several populations) to 8.79% (Alpechin population) in domesticated populations and from 1.65% (Ecuadorian population) to 70.57% (Malaysian population) in wild populations. In addition to the Malaysian population, the Far East Asian wild population also shows a high α value of 40.59%. These two extreme values are the result of a comparatively higher *ω_a_* and lower *ω_na_* (*Table 1*), and are in the range of what was measured in several animal species (Galtier, 2016). Accounting for the phylogenetic relationships of the populations, we find that these differences are marginally significant: domesticated populations have a lower α compared to wild populations (Generalised Least Square linear model, P value = 0.057 for non-Asian populations, 0.037 for Asian populations, see *Table 2*). We also find a marginal negative effect of the starter-based method, these populations having a lower α (P value = 0.075) compared to environmental populations. Repitching populations have no significant difference with the environmental ones (P value = 0.249). We further note a positive effect of *π_S_* on α (P value = 0.023). Including population-specific genes leads to more homogeneous results among populations, as all domesticated populations still have a very low α, while wild populations display higher values. Both α and *ω_a_* are significantly lower in domesticated vs wild populations, while *ω_na_* is significantly higher, as expected if domesticated populations have a lower effective population size. In this dataset, however, the remaining effect of population size (*π_N_*) is no longer significant after accounting for the life-history variables (Supplementary table 3).

**Table 1:** Genome-wide estimates of the rate of adaptive and non-adaptive non-synonymous substitutions in domesticated and wild populations of S. cerevisiae.

| LifeHistory | Method | Core genes |  |  | All genes |  |  | Core genes |  |  | All genes |  |  | Genome |
| --- | --- | --- | --- | --- | --- | --- | --- | --- | --- | --- | --- | --- | --- | --- |
| | | $\alpha$ | $\omega^a$ | $\omega^na$ | $\alpha$ | $\omega^a$ | $\omega^na$ | $\pi N$ | $\pi S$ | $\pi N/\pi S$ | $\pi N$ | $\pi S$ | $\pi N/\pi S$ | $\pi$ |
| Wild | Environmental | 2.68% | 0.004 | 0.142 | 54.58% | 0.0942 | 0.0783 | 4.75E-06 | 4.47E-06 | 1.06 | 1.73E-05 | 1.26E-04 | 0.14 | 8.49E-04 |
| Wild | Environmental | 70.57% | 0.105 | 0.044 | 74.35% | 0.1290 | 0.0445 | 4.25E-06 | 6.24E-05 | 0.07 | 1.44E-05 | 1.17E-04 | 0.12 | 1.97E-03 |
| Wild | Environmental | 40.59% | 0.059 | 0.087 | 52.56% | 0.0902 | 0.0813 | 1.13E-03 | 7.58E-03 | 0.15 | 1.20E-03 | 7.80E-03 | 0.15 | 4.26E-03 |
| Wild | Environmental | 1.65% | 0.002 | 0.143 | 52.45% | 0.0900 | 0.0816 | 6.69E-06 | 9.60E-06 | 0.70 | 1.61E-05 | 1.06E-04 | 0.15 | 1.50E-03 |
| Wild | Environmental | 14.71% | 0.022 | 0.125 | 25.89% | 0.0444 | 0.1271 | 1.39E-03 | 5.72E-03 | 0.24 | 1.47E-03 | 6.03E-03 | 0.24 | 3.52E-03 |
| Wild | Environmental | 10.07% | 0.015 | 0.132 | 26.74% | 0.0461 | 0.1260 | 8.18E-04 | 4.65E-03 | 0.18 | 8.58E-04 | 4.77E-03 | 0.18 | 2.60E-03 |
| Non-Asian & Domesticated | Environmental | 0.40% | 0.001 | 0.146 | 10.97% | 0.0189 | 0.1531 | 4.85E-04 | 1.86E-03 | 0.26 | 4.98E-04 | 1.92E-03 | 0.26 | 1.73E-03 |
| Non-Asian & Domesticated | Environmental | 8.79% | 0.013 | 0.136 | 27.35% | 0.0481 | 0.1278 | 7.06E-04 | 3.94E-03 | 0.18 | 8.10E-04 | 4.46E-03 | 0.18 | 3.76E-03 |
| Non-Asian & Domesticated | Repitching | 1.84% | 0.003 | 0.146 | 14.37% | 0.0250 | 0.1489 | 1.42E-03 | 6.64E-03 | 0.21 | 1.50E-03 | 7.01E-03 | 0.21 | 4.41E-03 |
| Non-Asian & Domesticated | Repitching | 0.00% | 0.000 | 0.149 | 0.03% | 0.0001 | 0.1748 | 6.33E-04 | 1.58E-03 | 0.40 | 6.67E-04 | 1.68E-03 | 0.40 | 1.52E-03 |
| Non-Asian & Domesticated | Repitching | 2.33% | 0.003 | 0.142 | 17.38% | 0.0297 | 0.1408 | 1.24E-03 | 4.78E-03 | 0.26 | 1.28E-03 | 4.91E-03 | 0.26 | 2.79E-03 |
| Non-Asian & Domesticated | Starter-based | 0.00% | 0.000 | 0.148 | 0.00% | 0.0000 | 0.1729 | 6.04E-04 | 1.46E-03 | 0.41 | 6.27E-04 | 1.47E-03 | 0.43 | 1.15E-03 |
| Non-Asian & Domesticated | Starter-based | 0.01% | 0.000 | 0.150 | 0.00% | 0.0000 | 0.1748 | 1.26E-03 | 3.55E-03 | 0.36 | 1.29E-03 | 3.68E-03 | 0.35 | 2.46E-03 |
| Non-Asian & Domesticated | Starter-based | 0.70% | 0.001 | 0.146 | 12.02% | 0.0207 | 0.1511 | 1.11E-03 | 4.99E-03 | 0.22 | 1.16E-03 | 5.24E-03 | 0.22 | 2.70E-03 |
| Non-Asian & Domesticated | Starter-based | 0.00% | 0.000 | 0.146 | 0.00% | 0.0000 | 0.1707 | 1.19E-03 | 3.22E-03 | 0.37 | 1.21E-03 | 3.25E-03 | 0.37 | 2.33E-03 |
| Asian & Domesticated | Environmental | 0.00% | 0.000 | 0.147 | 0.01% | 0.0000 | 0.1728 | 4.00E-04 | 1.30E-03 | 0.31 | 4.14E-04 | 1.31E-03 | 0.32 | 9.75E-04 |

**Table 2:**
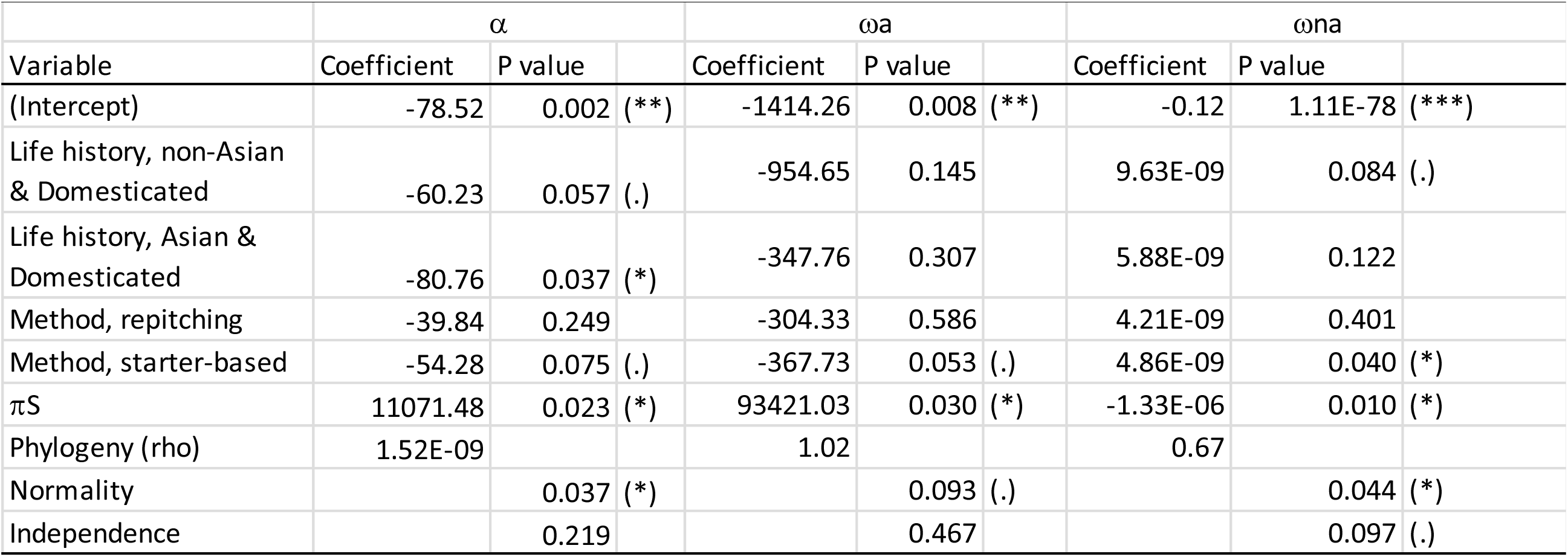
Effect of life history, culture method and effective population size (as estimated by *π*s) on the genome average rate of adaptive and non-adaptive non-synonymous substitutions.

As opposed to previous work based on the MK-test (Liti et al., 2009), and estimates from the closely-related wild yeast *S. paradoxus* (Gossmann et al., 2012), we found a positive rate of adaptive non-synonymous substitutions in several populations of *S. cerevisiae*. We hypothesise that the generally low rate of adaptive non-synonymous substitutions results from two possible, non-mutually exclusive causes: population size changes since the time of divergence, and intra genomic variation of the substitution rates. We will examine these two aspects in the following sections.

### *S. cerevisiae* population size decreased since the divergence with *S. paradoxus*

The historical change of population is an important factor shaping adaptive evolution (Charlesworth, 2009; Gossmann et al., 2012; Lanfear et al., 2014; M. Rousselle et al., 2018). We employed two approaches based on the sequentially Markov coalescent framework to infer the demographic histories of the selected populations. We first used the multiple sequentially Markov coalescent version 2, MSMC2 (Malaspinas et al., 2016) to get insights into populations sizes at the polymorphism scale (that is, from present time to the time of the most recent common ancestor of the sample). We then employed the IMCoalHMM approach to fit a model of isolation with migration between *S. cerevisiae* and *S. paradoxus* (Mailund et al., 2012), allowing us to estimate the population size at the time of the divergence between the two species. The estimated divergence time was found close to the most ancient time interval of the MSMC analysis, in agreement with the recent speciation between the two species (*Figure 2*). Estimating population sizes in number of individuals and determining coalescence events in number of years requires knowledge of the per-generation mutation rate and generation times, which are difficult to assert in the case of yeast, owing to the existence of an asexual phase. Assessing the impact of demography on the rate of adaptive substitutions, however, only requires information about the relative population sizes at the polymorphism and divergence level (Soni, Moutinho, & Eyre-Walker, 2022). Therefore, we report time in coalescence units and inferred historic variations of the inverse instantaneous coalescent rate (IICR), a measure proportional to *N_e_* when sequences are evolving neutrally (Chikhi et al., 2018; Mazet, Rodríguez, Grusea, Boitard, & Chikhi, 2016). We further conducted PSMC analyses on pairs of individuals in each population (*Figure 2*). This analysis showed no inconsistencies between individuals within each population, as could result, for instance, from population structure. This was also true for the Cocoa population, despite its paraphyly (*Figure 1*). As expected, increasing the sample size led to a higher resolution towards more recent times. All populations have a lower recent population size compared to the one of the *cerevisiae-paradoxus* ancestor. Furthermore, most populations have a decreasing population size. A notable exception is the North American Oak population, which shows a recent expansion, and the Mediterranean oak, which seems to have undergone a recent “burst” followed by another collapse.

**Figure 2:**
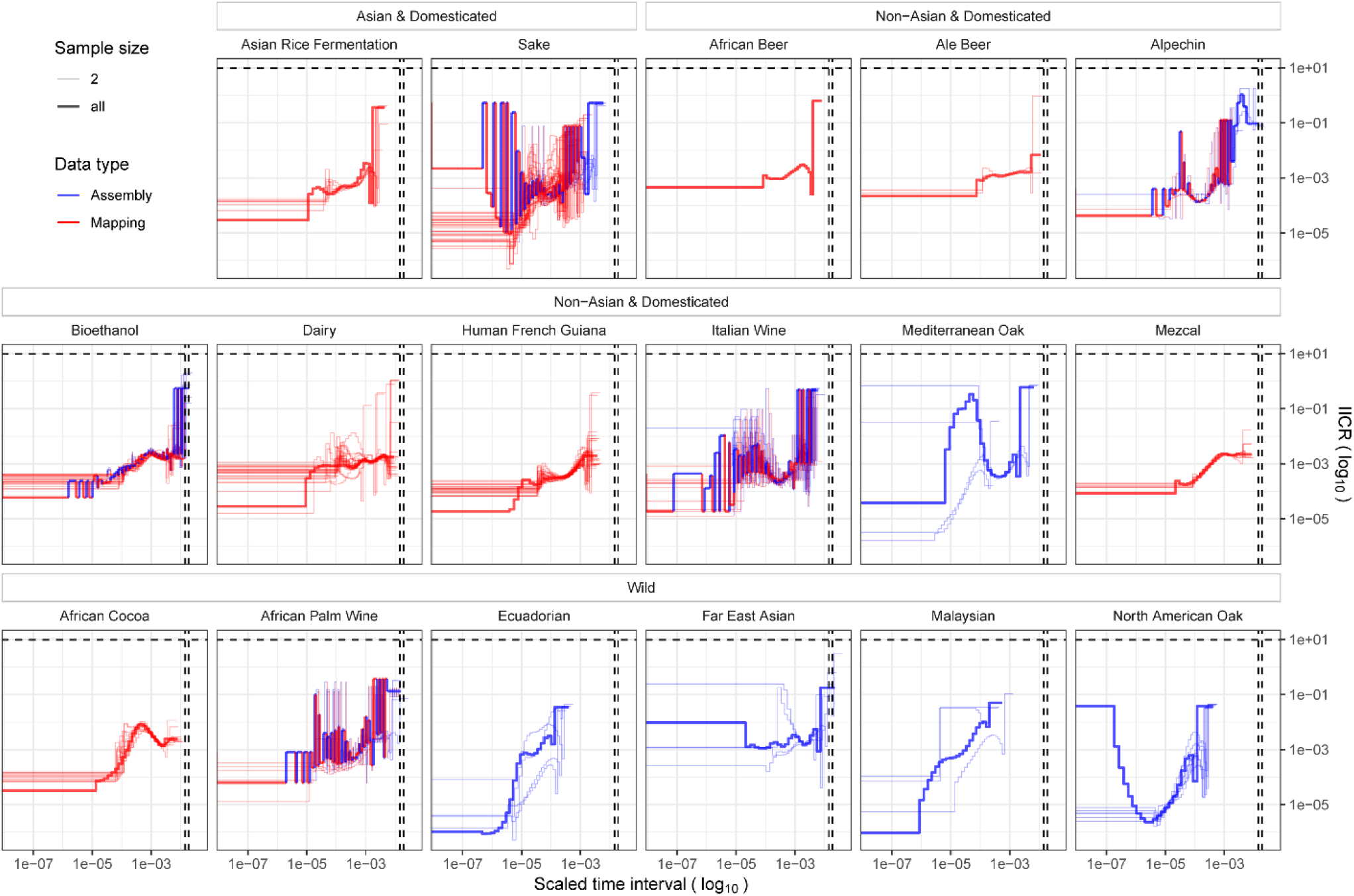
Demographic histories of *S. cerevisiae* populations. The inversed instantaneous coalescent rate (IICR, as inferred with MSMC2) is plotted as a function of time in coalescence units. The demographic inference was conducted either from *de novo* genome alignments (haploid or homozygous diploid isolates, blue lines) or from SNP calling after mapping reads to the *S. cerevisiae* reference genome (heterozygous diploid isolates, red lines). Thick lines represent the demography inferred from all isolates using MSMC. Thin lines represent the demography inferred from pairs of haplotypes using PSMC. Vertical dash lines: divergence time from *S. paradoxus*; horizontal dash lines: average ancestral IICR for the *S. paradoxus* – *S. cerevisiae* ancestor as estimated from IMCoalHMM

Linked selection, and in particular background selection (BgS), was shown to impact demographic inference (Boitard, Arredondo, Chikhi, & Mazet, 2022; Johri et al., 2021). Because the genome of *S. cerevisiae* is small and has a high density of coding regions, BgS can potentially be strong. We assessed the performance of MSMC2 in the presence of background selection in a *S. cerevisiae*- like genome by simulating data with the reference gene set and deleterious mutations (see Methods). We found that compared to simulations under a purely neutral scenario (*Figure 3A*), background selection led to an apparent recent increase in population size despite the real population size being constant (*Figure 3B*), recapping the results of previous studies (Boitard et al., 2022; Johri et al., 2021). When the real demography is declining, the resulting inferred demography shows a combination of decline followed by a recent expansion. In the light of these simulation results, we conclude that the observed decline of IICR in most yeast populations reflects a decrease in population size since the divergence with *S. paradoxus*. Yet any observed recent increase of the IICR might not reflect a population size recovery, as such an increase is expected under a scenario of constant or decreasing population size with BgS.

**Figure 3:**
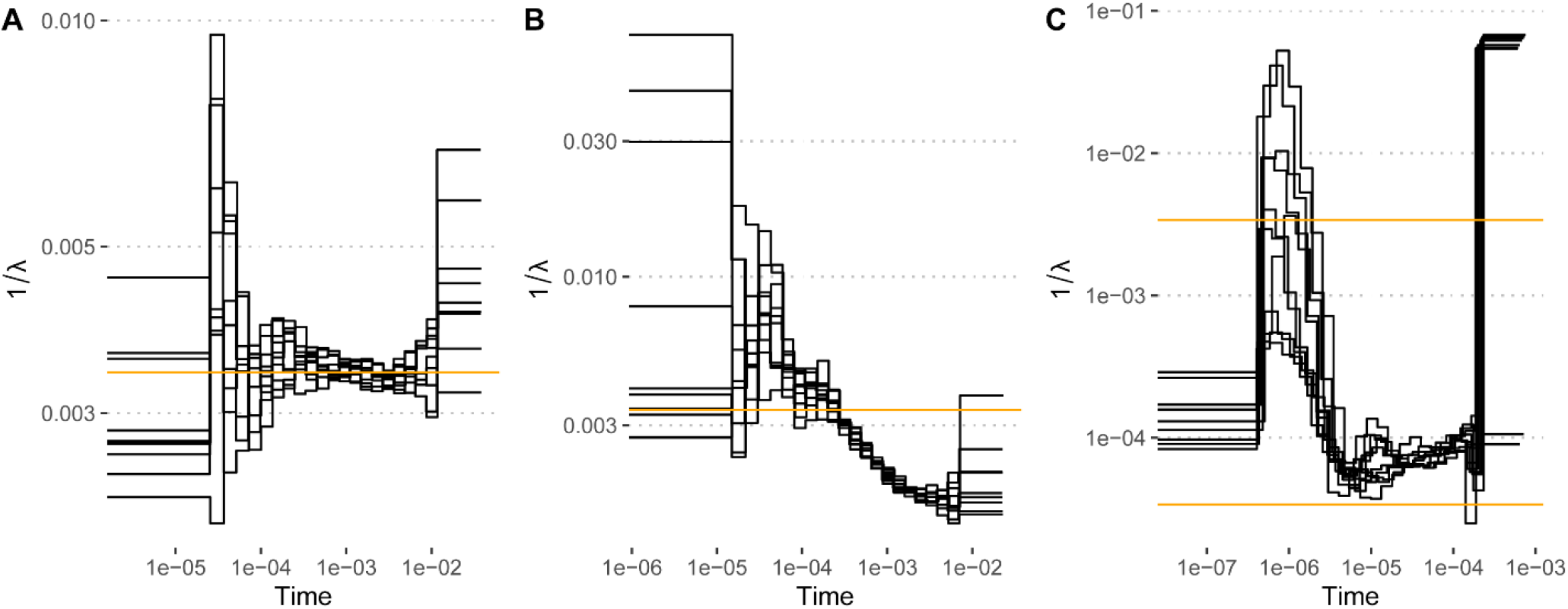
Demography recovery from simulated data with yeast genome characteristics, including chromosome sizes, gene content and callability mask. Ten replicates are plotted with black lines. A) Simulations under a neutral scenario and a flat demography (constant population size, plotted in orange). B) Simulations under a scenario with background selection and a flat demography (constant population size, plotted in orange). C) Simulations under a scenario with background selection and an exponentially declining population (from 10,000 individuals, upper orange line) to 100 individuals (lower orange line).

Historic demographic changes are a known confounding factor in inferring adaptive evolution, as fluctuations in population size can influence segregating frequencies of polymorphisms independent, or even in spite, of selection acting on sites (Olson-Manning, Wagner, & Mitchell Olds, 2012; M. Rousselle et al., 2018; Soni et al., 2022). In particular, decreasing population sizes since the population divergence leads to underestimations of the rate of adaptive non-synonymous substitutions, an argument that was previously invoked to explain the low rate of adaptive non synonymous substitutions in Primates and could explain the generally low rate of adaptive mutations in yeast too (Soni et al., 2022). Moreover, a lower present effective population size may lead to artificial positive correlations between the rate of adaptive substitutions and any variable that is itself positively correlated with the strength of negative selection. In the following, we estimate the rate of adaptive non-synonymous substitutions (*ω_a_*) in all yeast populations and assess the impact of intra-genomic factors on its distribution. Because of the difference in population sizes, we further control for the strength of negative selection.

### Domestication-associated gene sets harbour lower rates of non-adaptive evolution

To gain insights into adaptive evolution in wild and domesticated populations of *S. cerevisiae*, and in particular niche-adaptation of domesticated populations of *S. cerevisiae*, we focused on genes involved in known cellular processes that are implicated in the domesticated use of *S. cerevisiae*. Specifically, domesticated yeasts largely abandon meiosis and are exposed to strong osmotic stresses due to high concentrations of mono and disaccharides such as maltose, xylose, glucose, sucrose and fructose present early-on in fermentation reactions (Betlej et al., 2020; De Chiara et al., 2022), and also use these domestication-associated carbon sources more efficiently (De Chiara et al., 2022). Furthermore, domesticated isolates are oftentimes exposed to toxic metals, such as those present in pesticides, including copper and arsenic (Bencko & Yan Li Foong, 2017; Pietrzak & McPhail, 2004; Pontes, Čadež, Gonçalves, & Sampaio, 2019). These exposures are all typical of domestication-associated environments due to the processes involved in industrial fermentation and agriculture (Legras et al., 2018). Therefore, we expect to detect signatures of relaxed selection on meiosis-associated genes, while for osmotic and metal stress-associated genes, we expect an increased rate of adaptive non-synonymous substitutions in domesticated populations over the wild populations. Conversely, wild populations are more tolerant to natural stresses such as heat stress and oxidative stress compared to domesticated populations (De Chiara et al., 2022; Gallone et al., 2016). From these different environmental exposures and life cycle characteristics, we have compiled seven gene sets. We will refer to these resulting gene sets as “Meiosis” (N = 125), “Osmotic Stress” (N = 17), “Domestic Carbon” (N = 81), “Metal Stress” (N = 103), “Heat Stress” (N = 25), and “Oxidative Stress” (N = 48), respectively. Genes that were found to be annotated in several of these processes were combined into a category labelled “Multiple” (N = 59), and genes not annotated in any of these processes were categorised under the label “Other genes” (N =2,739). We hypothesise that genes involved in stress responses and carbon metabolism have a higher rate of adaptive non-synonymous substitutions in domesticated populations, while genes involved in meiotic processes should display signature of relaxed purifying selection, with a higher rate of non-adaptive non-synonymous substitutions.

Linear models showed that the gene set variable only had an impact on *ω_na_*; that is, the rate of adaptive substitutions did not differ significantly between the gene categories (*Figure 4 and Table 3*). Conversely, several gene categories had a significantly lower *ω_na_* compared to unlabelled genes: “metal stress”, “domestic carbon”, “osmotic stress”, “oxidative stress”, as well as genes with multiple labels. These results indicate that the corresponding genes undergo stronger purifying selection. However, this effect was found to be independent from life history or culture method (no interaction term was found significant), suggesting that this feature is general to *S. cerevisiae* and not affected by domestication. Domestication was found to have a significant positive effect on *ω_na_* and a negative effect on *ω_a_*, although the latter should be taken with caution as (1) the model residues did not fulfil the independence and normality assumption, and (2) the effect vanished when only strictly positive *ω_a_*were considered (*Table 3*). Repitching and starter based populations had a significantly lower *ω_a_* compared to environmental populations. They also had a higher *ω_na_*, although the effect vanished when considering only strictly positive *ω_a_*. Finally, *π_N_* had a positive effect on *ω_a_*, and a negative one on *ω_na_*, both weakly significant, consistent with the hypothesis that population with larger effective size have more efficient negative and positive selection.

**Figure 4:**
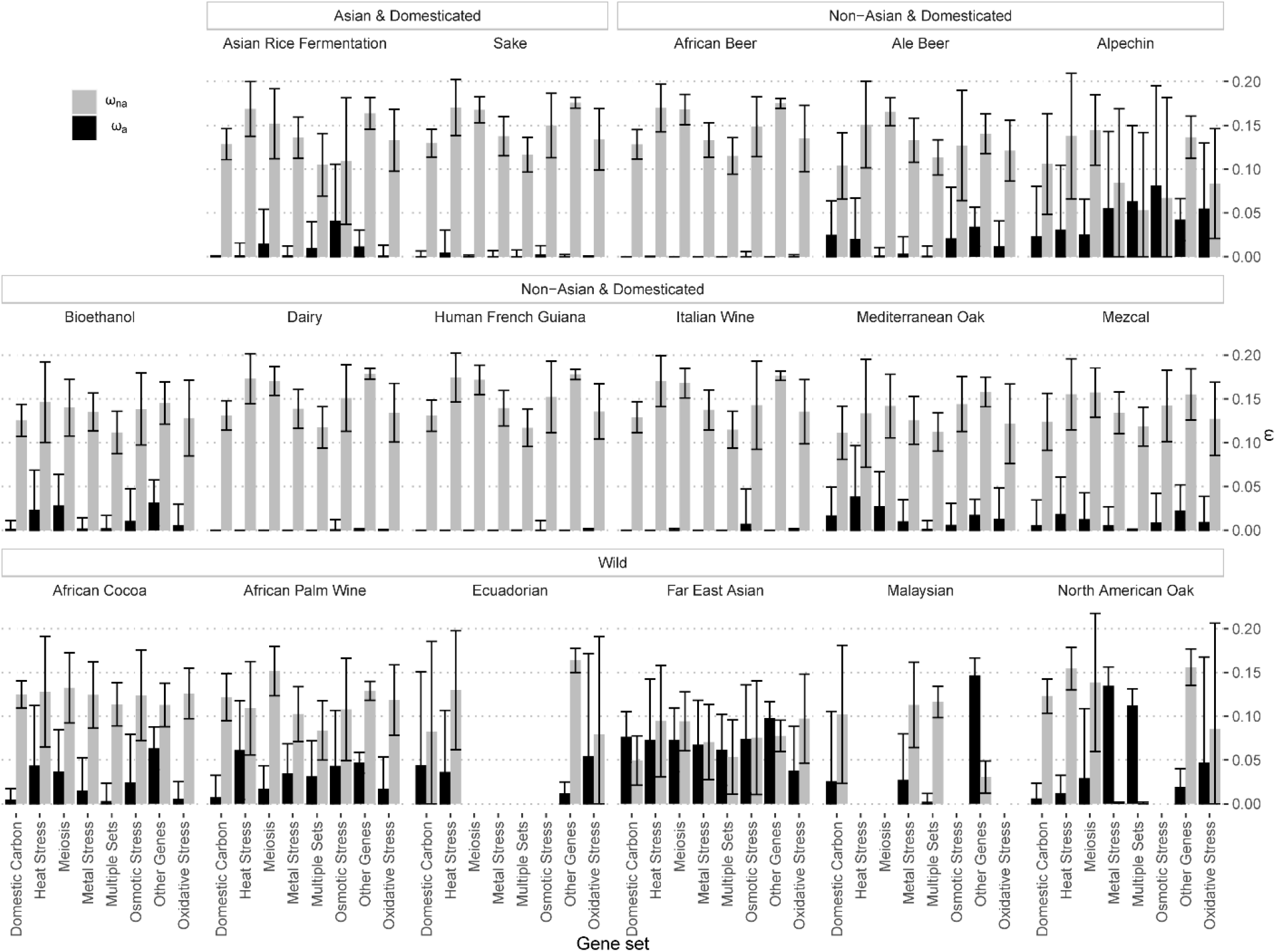
Rates of adaptive (ω_a_) and non-adaptive (ω_na_) non-synonymous substitutions in distinct gene set. Bars indicate the mean over a 150 bootstrap replicates, and the error bars the corresponding 95% confidence interval (1.96 * standard deviation).

**Table 3:**
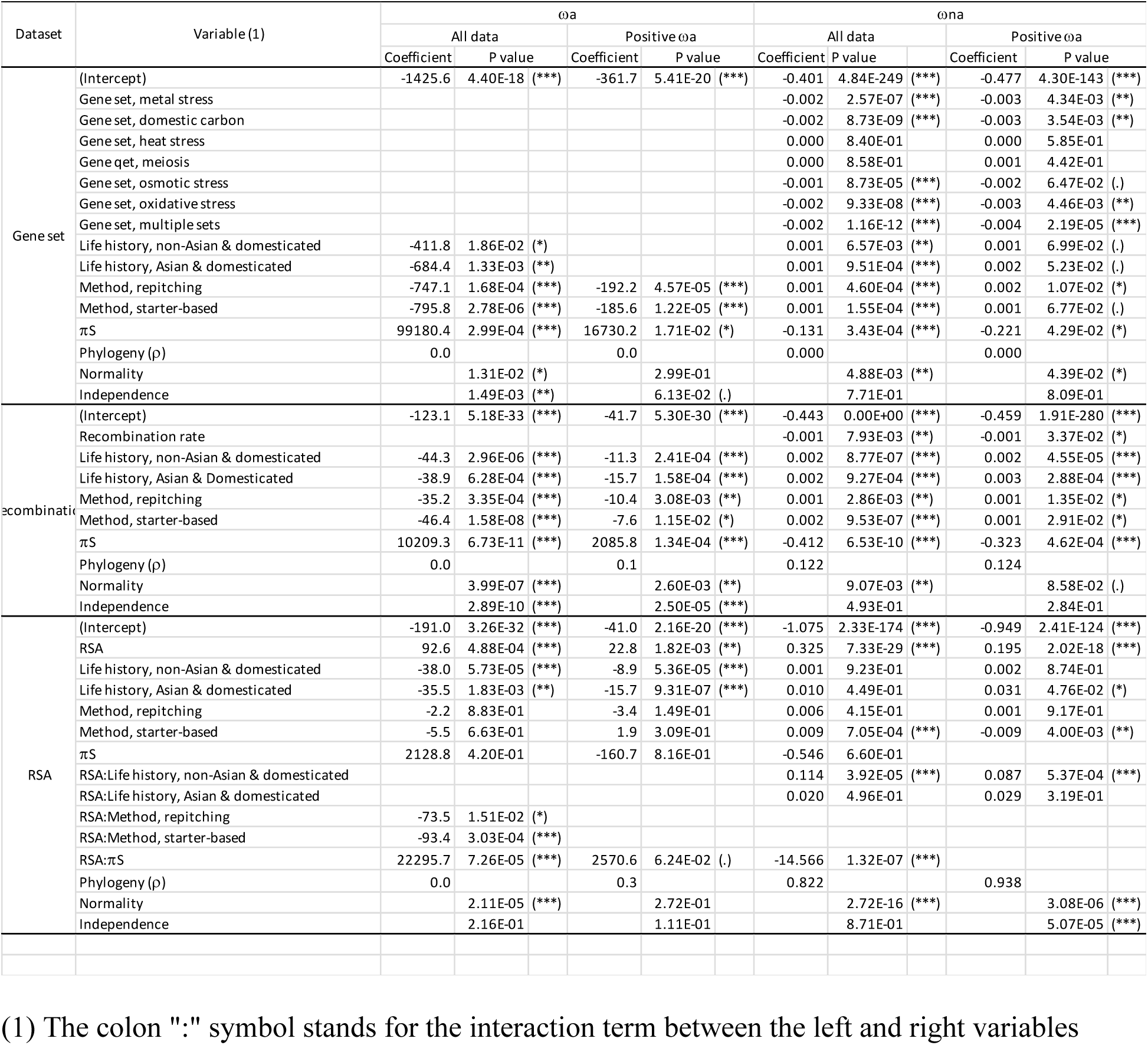
Effect of the gene functional category, local recombination rate and solvent exposure on the rate of adaptive and non-adaptive non-synonymous substitutions. Generalised least square models accounting for populations’ historical relationships. The reported models are the most parsimonious models according to Akaike’s information criterion, after transforming the response variable with a Box-Cox transform. Variable with an empty coefficient were not retained in the final model, for a given response variable.

This gene set analysis did not allow to confirm *a priori* defined gene categories as key players in the adaptive process linked to domestication. Their observed higher level of conservation is more likely to be the result of their central functions than the consequence of domestication. Nonetheless, these results suggest that domesticated populations undergo a lower rate of adaptive - and a higher rate of non-adaptive - non-synonymous substitution, and that effective population size, as measured by *π_N_*, affects both rates.

### No apparent correlation between the recombination rate and the rate of adaptive non synonymous substitutions in core yeast genes

We sought to assess how the non-synonymous substitution rates varied along the genome of *S. cerevisiae*. Recombination may affect the substitution rate at positions linked to loci under selection, a mechanism termed linked selection. A lower recombination rate is expected to slow down the fixation of advantageous mutations and reduce *ω_a_*, but also to reduce the efficacy of purifying selection and lead to an increased fixation of deleterious mutations, thus increasing *ω_na_*. To investigate the role of recombination in shaping the rate of adaptive evolution across genes in our *S. cerevisiae* populations, we binned genes into categories of approximately equal sizes according to the broad recombination landscape of *S. cerevisiae* (Liu et al., 2018) and quantified the rate of adaptive evolution across categories. We found no relation between the crossing over rate and the rate of adaptive evolution in *S. cerevisiae*, and a weak but significant negative relationship with *ω_na_* (*Figure 5 and Table 3*), suggesting that regions with a higher recombination rate tend to have a more efficient purifying selection and fix less deleterious mutations. The absence of a relation between *ω_a_* and the recombination rate indicates a limited role of Hill-Robertson interference in determining the rate of adaptive evolution in yeast. This contrasts with organisms like *Drosophila melanogaster*, where the recombination rate was shown to have a strong positive effect on the rate of adaptive evolution (Castellano, Coronado-Zamora, Campos, Barbadilla, & Eyre-Walker, 2016), but also *Zymoseptoria tritici*, a fungal pathogen of wheat (Grandaubert et al., 2019). We further note that, as with whole genome estimates of *ω_a_* and *ω_na_*, the effect of life history and culture method were significant: domesticated populations have a lower rate of adaptive and a higher rate of non-adaptive non-synonymous substitutions. Population size (as estimated by size *π_N_*) had a significant positive effect on *ω_a_*, and a significant negative effect on *ω_na_*, indicating that both positive and negative selection was more efficient in larger populations. We note that, despite variable transformation, the model fit was relatively poor, with significant departure from normality and independence of residues, indicating that the P-values of the model coefficients may be biased (Table 3). Furthermore, we note that the recombination rate is negatively correlated with codon usage bias, which points at the eventuality that codon usage bias masks the effect of recombination; a possibility that we investigate in more detail below.

**Figure 5:**
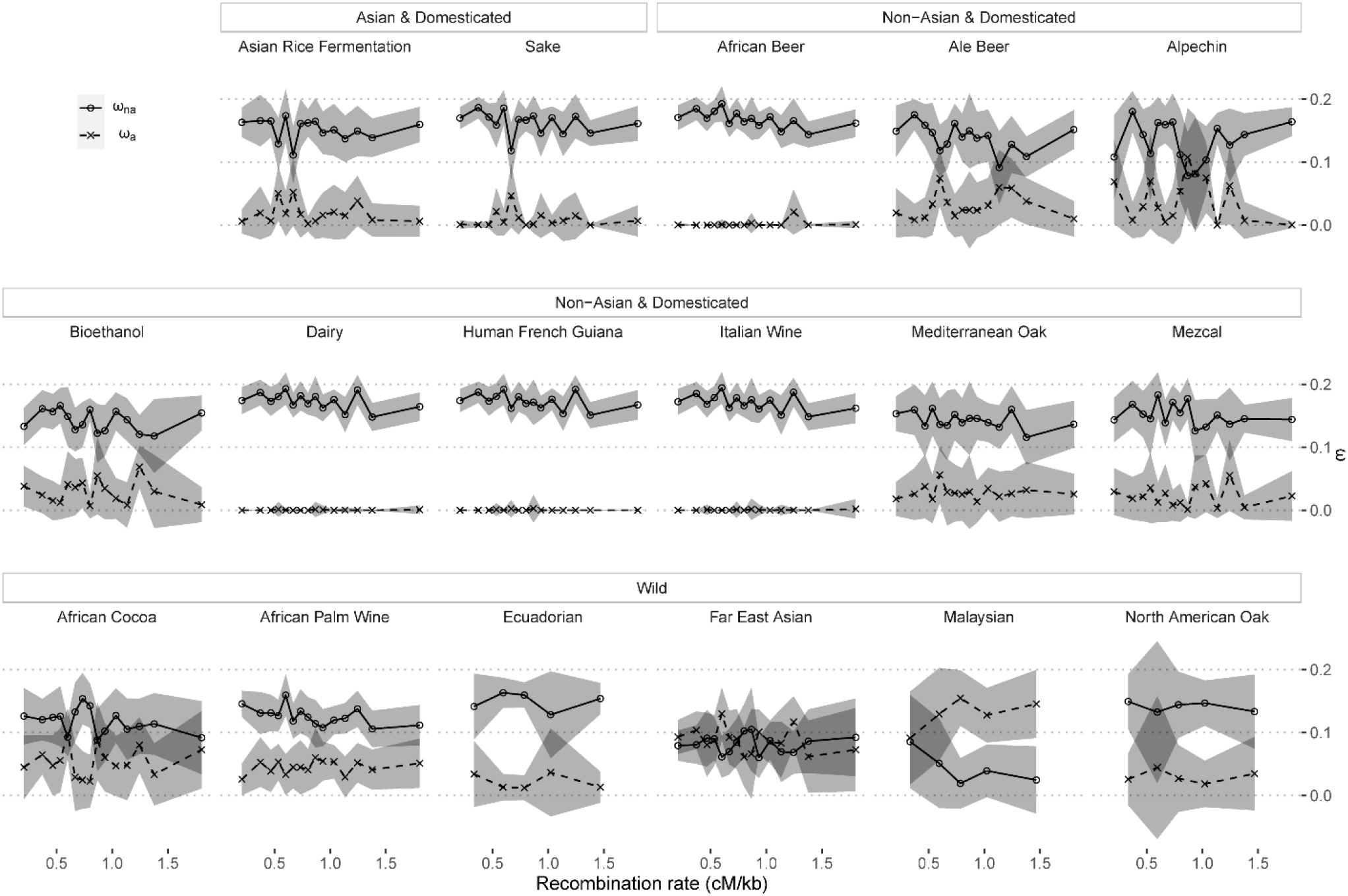
Rates of adaptive (ω_a_) and non-adaptive (ω_na_) non-synonymous substitutions in regions with distinct recombination rates. Points and lines indicate the mean over a 150 bootstrap replicates, and the shaded area the corresponding 95% confidence interval (1.96 * standard deviation).

### Protein structure shapes the rate of adaptive evolution within genes

Lastly, we assessed the impact of the residues solvent exposure, as it was recently shown to be a major determinant of substitution rates in *Arabidopsis thaliana*, *Drosophila melanogaster* (Moutinho et al., 2019), and Primates (Soni et al., 2022), with adaptation predominantly occurring at solvent-exposed residues in these species. To gain a more in-depth understanding of adaptive evolution at the protein-structural level in domesticated and wild populations of *S. cerevisiae*, we computed the rate of adaptive non-synonymous substitutions in residue categories with different levels of relative solvent accessibility (RSA) and inferred *ω_a_* and *ω_na_* in these categories. We found that RSA had a strong positive, significant effect on both rates, exposed residues having a higher rate of adaptive and non-adaptive substitutions (*Figure 6* and *Table 3*). As before, we find that domesticated populations had a significantly lower *ω_a_*, but culture methods had a significant interaction with RSA (the negative effect was stronger at exposed residues). The effect of population size (*π_N_*) was also significant as an interaction term with RSA: larger populations tend to undergo more adaptive substitutions at exposed residues.

**Figure 6:**
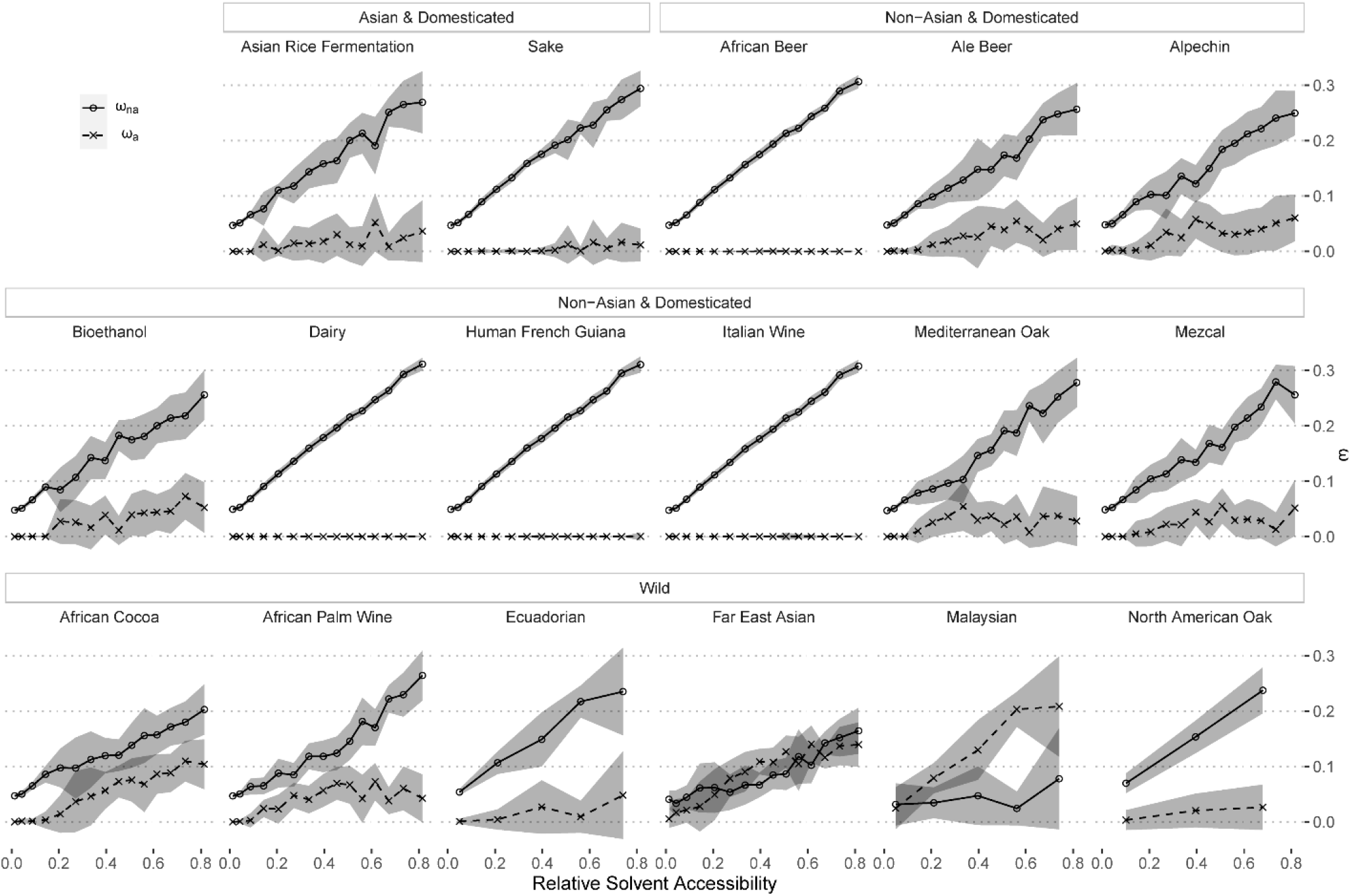
Rates of adaptive (ω_a_) and non-adaptive (ω_na_) non-synonymous substitutions in sites with distinct relative solvent accessibility (RSA). Points and lines indicate the mean over a 150 bootstrap replicates, and the shaded area the corresponding 95% confidence interval (1.96 * standard deviation).

We investigated whether the correlation between *ω_a_* and RSA could be an artefact of the decreasing population size (Soni et al., 2022). Such a correlation may artificially occur if RSA was positively correlated with the strength of purifying selection. We note, however, that RSA is positively correlated with *ω_na_*, implying that the strength of selection is negatively correlated with RSA. Demography is expected to bias the correlation in the opposite direction of the observed one, which is, therefore, genuine, similar to what was reported in Primates (Soni et al., 2022).

### The impact of codon usage bias on the observed patterns of adaptive evolution

Finally, we assessed whether selection on codon usage could bias our estimates of adaptive rates. The Mac-Donald-Kreitman formalism involves the use of a neutral reference, for which synonymous mutations are used. Selection for optimal codons on synonymous mutations may, therefore, affect the estimates of adaptive substitution rates. As we are investigating differences in rates between genes and populations, we assessed whether differences in codon usage could explain the correlations we observed, or on the contrary, hide an existing correlation signal. We computed codon usage biases for all genes and populations of our dataset, and we found no evidence for distinct average codon usage between populations (pairwise Kendall’s rank correlation tests on relative synonymous codon usage (RSCU) values, minimum correlation > 0.999, maximum P value < 8.408e-131). Pulling all populations together (with the exclusion of Ecuadorian, Malaysian, and North American Oak, for which 15 recombination rates and RSA rates could not be computed), we did not find any difference in codon usage between gene sets (Kruskal Wallis test, P value = 0.8827). Conversely, we find a significant effect of the recombination rate on codon usage bias (Kendall’s tau = −0.85, P Value = 4.073e-7, correlation between the crossing over rate and the mean deviation of RSCU from 1 for each recombination rate category, Supplementary Figure 3), suggesting that genes in low recombination rates exhibit strong codon bias than gene in highly recombining regions. An effect of recombination on the efficacy of selection would lead to the opposite pattern, where highly recombining regions should display stronger codon bias. Therefore, the observed correlation between codon usage bias and recombination rate is probably spurious, resulting from a covariate such as gene expression level. To test whether codon usage bias could mask the effect of recombination on the rate of adaptive non-synonymous substitutions, we re-estimated *ω_a_* and *ω_na_* in all recombination categories, using the synonymous site frequency spectrum of the highest recombination rate category, as it is the least affected by selection on codon usage. As a result, the crossing over rate has a marginally significant positive effect on the rate of adaptive non-synonymous substitutions (P value = 6.00e- 2, Table 4) and a negative effect on the rate of non-adaptive non-synonymous substitutions (P value = 2.33e-3). Domesticated isolates have a significantly lower *ω_a_* and a significantly higher *ω_na_*, while the method (repitching vs starter-based), as well as the effective population size (*π_N_*), have no significant effect. In conclusion, correcting for codon usage bias revealed a small effect of the recombination rate, the direction of which is consistent with the action of linked selection. The effect of the domestication variable remains strong after the correction, in contrast to the inoculation method and population size, whose effect proved to be not robust.

**Table 4:** Effect of local recombination rate and solvent exposure on the rate of adaptive and non-adaptive non-synonymous substitutions after correction for selection on synonymous codon usage.

| Dataset | Variable | $\omega_a$ | | | | $\omega_{na}$ | | | |
| --- | --- | --- | --- | --- | --- | --- | --- | --- | --- |
| | | All data | | Positive $\omega_a$ | | All data | | Positive $\omega_{na}$ | |
|  |  | Coefficient | P value | Coefficient | P value | Coefficient | P value | Coefficient | P value |
| recombinatic | (Intercept) | -1223.4 | 3.05E-12 (***) | -253.2 | 5.87E-14 (***) | -0.394 | 0.00E+00 (***) | -0.487 | 1.17E-245 (***) |
|  | Rec | 337.3 | 4.21E-02 (*) | 64.7 | 3.81E-02 (*) | -0.001 | 2.21E-03 (**) | -0.003 | 1.51E-03 (**) |
|  | Life history | -1235.9 | 1.27E-22 (***) | -185.1 | 3.99E-14 (***) | 0.002 | 4.64E-13 (***) | 0.005 | 1.98E-11 (***) |
|  | Life history | -1196.2 | 6.80E-14 (***) | -204.9 | 1.40E-10 (***) | 0.002 | 1.79E-11 (***) | 0.005 | 1.48E-08 (***) |
|  | Method, repitching |  |  |  |  | 0.000 | 8.17E-01 |  |  |
|  | Method, starter-based |  |  |  |  | 0.000 | 5.42E-02 (.) |  |  |
|  | Phylogeny | 0.2 |  | 0.2 |  | 0.466 |  | 0.409 | NA |
|  | Normality |  | 3.25E-07 (***) |  | 1.30E-01 |  | 1.63E-01 |  | 2.09E-01 |
|  | Independence |  | 5.31E-04 (***) |  | 8.69E-02 (.) |  | 9.91E-04 (***) |  | 3.90E-02 (*) |
| RSA | (Intercept) | -1713.6 | 9.72E-37 (***) | -158.4 | 9.67E-35 (***) | -1.043 | 5.66E-190 (***) | -0.905 | 9.50E-117 (***) |
|  | RSA | 1082.4 | 7.99E-07 (***) | 145.3 | 2.23E-17 (***) | 0.319 | 6.40E-38 (***) | 0.178 | 1.07E-19 (***) |
|  | Life history | -385.3 | 6.72E-07 (***) | -8.3 | 4.79E-01 | -0.003 | 7.84E-01 | -0.008 | 5.24E-01 |
|  | Life history | -213.0 | 2.09E-02 (*) | -5.8 | 7.69E-01 | -0.002 | 8.54E-01 | -0.013 | 3.25E-01 |
|  | Method, re | 168.1 | 1.73E-01 | -8.0 | 2.99E-01 | -0.001 | 9.21E-01 |  |  |
|  | Method, st | 139.4 | 1.81E-01 | 8.3 | 2.65E-01 | -0.002 | 6.50E-01 |  |  |
| | $\pi_S$ | 11740.2 | 5.86E-01 | 3705.1 | 1.28E-03 (**) | 0.073 | 9.34E-01 | -2.449 | 5.43E-03 (**) |
|  | RSA:Life history, non-Asian & domesticated |  |  | -50.8 | 2.13E-02 (*) | 0.104 | 7.88E-05 (***) | 0.093 | 3.02E-04 (***) |
|  | RSA:Life history Asian & domesticated |  |  | -74.8 | 2.22E-02 (*) | -0.002 | 9.24E-01 | 0.052 | 3.93E-02 (*) |
|  | RSA:Metho | -981.8 | 7.93E-05 (***) |  |  | 0.016 | 4.83E-01 |  |  |
|  | RSA:Metho | -1018.1 | 1.83E-06 (***) |  |  | 0.015 | 5.16E-02 (.) |  |  |
| | RSA:$\pi_S$ | 201172.2 | 1.25E-05 (***) | | | -16.375 | 1.67E-15 (***) | | |
|  | Phylogeny | 0.0 |  | 0.0 |  | 0.949 |  | 0.854 |  |
|  | Normality |  | 2.87E-03 (**) |  | 9.95E-01 |  | 3.72E-16 (***) |  | 1.29E-06 (***) |
|  | Independence |  | 5.91E-01 |  | 5.64E-02 (.) |  | 2.03E-01 |  | 3.21E-01 |

Finally, looking at codon usage bias per RSA category, we find that the first axis of a within-class correspondence analysis was well described by RSA (*Supplementary Figure 4A*) and that the mean codon usage bias was significantly negatively correlated with the average RSA (*Supplementary Figure 4B*, Kendall’s tau = −0.87, -value = 1.543e-07, see Methods). A strong relation between RSA and codon usage has been previously reported in several species (Ramsey, Scherrer, Zhou, & Wilke, 2011; Zhou, Weems, & Wilke, 2009), suggesting that synonymous codon usage is under selection in a structure-dependent manner. As the assumption of neutrality of synonymous sites is central in our DFE-based adaptive protein evolution inference methods, we reanalysed the relation between RSA and the rate of adaptive evolution using synonymous sites only from the most exposed RSA category, as these showed the least amount of codon bias. Overall, the results from this reanalysis are strongly consistent with our initial findings (*Table 4*), confirming that the relation between RSA and the rate of adaptive evolution is independent of the stronger codon bias in buried residues.

## Discussion

The rate of molecular adaptive evolution was previously reported to be low in *Saccharomyces*, with a rate of adaptive non-synonymous substitutions found to be equal to zero (Gossmann et al., 2012; Liti et al., 2009). This low rate contrasts with the high rate of molecular evolution of other fungal species, in particular fungal pathogens (Grandaubert et al., 2019; Schweizer et al., 2021). Here, we showed that despite this average low rate, signatures of adaptation are present in *S. cerevisiae* populations, most notably in wild populations. We further determined which factors underlie the variation of substitution rates between populations and within genomes, using linear models accounting for the historical relationships between populations. Our phylogenetic models generally reveal a stronger phylogenetic component (parameter ρ) for the rate of non-adaptive non synonymous substitutions compared to the rate of adaptive non-synonymous substitutions, indicating that adaptive substitutions occur in a population-specific manner.

To search for regions with signature of adaptation, we assessed three factors that may impact the substitution rate: the gene functional category, the local recombination rate, and, at the site level, the relative solvent accessibility. While all these factors proved to have a significant impact on the rate of non-adaptive non-synonymous substitutions, *ω_na_*, only solvent accessibility significantly and consistently impacted the rate of adaptive non-synonymous substitutions, *ω_a_*. This effect of RSA is consistent with previous reports from *Drosophila*, *Arabidopsis* (Moutinho et al., 2019) and human (Soni et al., 2022) emphasizing its generality across organisms with distinct life-history traits. The impact of RSA on *ω_na_* can be attributed to mutations having a generally less deleterious effect; exposed residues having on average less contact with other residues and, therefore, having more flexibility to cope with a sidechain change. We further showed that the correlation of *ω_a_* with RSA was not an artefact of its correlation with the strength of purifying selection.

We estimated substitution rates in different gene categories, based on the possible roles the underlying genes may play in the domestication process. However, we did not find any significant differences in *ω_a_* between these gene classes. Our results revealed that domestication is associated with a general reduction of the rate of adaptive substitutions, both in core and dispensable genes, and are consistent with domesticated populations having undergone a reduction in their effective population size, limiting the rate of adaptive – and increasing the rate of non-adaptive – non synonymous substitutions. While these results do not preclude the occurrence of some adaptive substitutions, it parallels previous reports of gene copy number variations (CNV) associated with domesticated populations (Duan et al., 2018), supporting CNV mutations as a driver of the molecular changes underlying the domestication process. Furthermore, the comparatively constant environment associated with human practices could temper the rate of beneficial mutations. We note that our analysis focused on the genes recovered from all populations, and thus did not assess molecular rates in population-specific genes. Assessing the substitution rate in population-specific duplicated genes offers a promising perspective to dissect the genetic mechanisms of yeast domestication.

While demography inference in all populations revealed a general trend of decreasing population sizes, we did not observe systematic differences between domesticated and wild populations. Historical effective population size, however, as estimated by the neutral diversity *π_S_* proved to be a significant determinant of the rates of both adaptive and non-adaptive substitutions. These results, therefore, suggest that in *S. cerevisiae* populations, adaptation is limited by the supply of beneficial mutations, as observed is several animal taxonomic groups (Marjolaine Rousselle et al., 2020).

Finally, we investigated the evidence for signatures of linked selection on the substitution rate. Using a previously published estimated map of crossing-over rates along the genome of *S. cerevisiae*, we found no correlation between the rate of adaptive substitutions and the recombination rate. We concluded that adaptive evolution in *S. cerevisiae* is only weakly, if at all, impeded by Hill-Robinson interference (Comeron, Williford, & Kliman, 2008; Hill & Robertson, 1966). This result is most likely explained by the combination of a globally low rate of adaptive substitutions and a high recombination rate along the *S. cerevisiae* genome. Conversely, we found evidence for the occurrence of background selection (BgS), as we observed a significant negative correlation between the recombination rate and the rate of non-adaptive non-synonymous substitutions. BgS is expected to be high in *S. cerevisiae*, given its high gene density (more than 73% of the genome encodes an exon). Although significant, the correlation between the recombination rate and *ω_na_* was found to be low, probably because of the high recombination rate resulting in a low linkage disequilibrium.

In summary, our analyses revealed the occurrence of adaptive non-synonymous substitutions in several populations of *S. cerevisiae*. While we found the rate of adaptive substitutions to be generally low, we showed that accounting for intra and inter-genic factors unravelled signatures of molecular adaptation. Adaptive substitutions were notably found in population specific genes, and otherwise at the solvent-exposed surface of proteins, following a general trend observed is other species of animals and plants and suggesting that these mutations were not directly linked to the domestication process. Moreover, we showed that domestication is associated with a reduction of the rate of adaptive non-synonymous substitutions, and that in *S. cerevisiae* wild and domesticated populations, adaptation is generally limited by the supply of mutations.

## Acknowledgements

We would like to thank Ana Filipa Moutinho, Adam Eyre-Walker, and Nicolas Galtier for discussions on this project. We thank Ana Filipa Moutinho for help with running the NetSurfP-2.0 software and Haozuan Liu and Jianzhi Zhang for kindly providing data on recombination rates across the *S. cerevisiae* genome, and Fernanda Trancoso, Ana Filipa Moutinho, and Chrats Melkonian for critical reading of the manuscript. J. Y. D. would like to thank Lucie Vivier, wine grower and winemaker in Charente, France, for her explanations on yeast industrial usage in winemaking. J.Y.D. acknowledges funding from the Max Planck Society.

## Data availability

All data and scripts are accessible via FigShare at http://doi.org/10.6084/m9.figshare.21725513 (Raas & Dutheil, 2022)

## Author Contributions

MWDR: Conceptualization, Formal analysis, Investigation, Data curation, Writing – Original draft, Writing – Review & Editing, Visualization

JYD: Conceptualization, Validation, Formal analysis, Investigation, Writing – Review & Editing, Visualization, Supervision

## Supplementary tables

**Supplementary Table 1**: List of all selected isolates, their designated population, assembly identifiers, zygosity, ploidy and aneuploidy status.

**Supplementary Table 2:** Length of nucleotide alignment for each population after whole genome alignment and after extraction of CDS regions, and the total number of genes recovered from each population.

**Supplementary Table 3**: Effect of life history, culture method and effective population size (as estimated by *π*s) on the genome average rate of adaptive and non-adaptive non-synonymous substitutions when accounting for population-specific genes.

## Supplementary figures

**Supplementary Figure 1: Demography recovery from simulated data with yeast genome characteristics, under a neutral scenario and flat demography.** The true demography (constant population size) is plotted in orange. The inferred demography from ten replicates is plotted with black lines. A) Demography inference using MSCM2’s default time discretization. B) Demography inference using a 1*5+20*1+1*5 time discretization scheme. C) Demography inference using a 1*5+20*1+1*5 time discretization scheme and callability mask of the real Far East Asia population.

**Supplementary Figure 2: UpSet plot showing the distribution and overlap of genes across different populations.** Diagram summarising the number of genes for all unique intersection set. Corresponding intersection sets are represented with the dots connected by lines in the plot underneath the main bar plot, where filled-in dots signify a population is part of the set, and where a non-included population is shown as a light grey dot with possibly the dark line spanning across it. The set size plotted on the left of the figure represents the total amount of genes recovered from each of the populations. This figure was created using the ‘UpSetR’ package v.1.4.0 (Conway, Lex, & Gehlenborg, 2017).

**Supplementary Figure 3: Codon usage analysis with respect to the crossing-over rate.** A) within-group correspondence analysis. Labels correspond to the 15 categories of sites with increasing recombination rate. B) Codon usage bias as a function of recombination rate class. Codon usage bias was measured as the mean relative synonymous codon usage (RSCU) – 1 (absolute value).

**Supplementary Figure 4: Codon usage analysis with respect to relative solvent accessibility (RSA).** A) within-group correspondence analysis. Labels correspond to the 15 categories of sites with increasing RSA. B) Codon usage bias as a function of RSA class. Codon usage bias was measured as the mean relative synonymous codon usage (RSCU) – 1 (absolute value).

