## Supplementary materials for "The rate of adaptive molecular evolution in wild and domesticated *Saccharomyces cerevisiae* populations"

#### Table of Contents:


| | |
|---|---|
| **Supplementary Methods** | Page 2 |
| **Supplementary Results** | Page 4 |
| **Supplementary Tables** | Page 6 |
| **Supplementary Figures** | Page 8 |
| **References** | Page 14 |

### Supplementary Methods

#### Genome-wide average population recombination rates

We used iSMC v0.0.23 to get a rough estimate of the genome-wide recombination rate average of each population (Barroso, Puzović, & Dutheil, 2019). We used either haploid or homozygous diploid isolates. We exported the corresponding filtered genome alignment to Fasta files (one per chromosome), filling uncalled positions with 'N' characters. Chromosome alignments were then concatenated into a single alignment and breakpoints were recorded in order to generate iSMC's 'tab' files. A standard model with homogeneous recombination rate was used. The corresponding estimates are provided in *Supplementary Table 4*.

#### Simulations

We conducted simulations with characteristics similar to those of our data set to assess the power of demography inference methods with yeast genome data. We first simulated ten replicates of a 10,000 individuals population evolving under a constant demography. Simulations were conducted using the 'msprime' software v1.1.0 (Baumdicker et al., 2021). A single chromosome of size 12,071,326 bp was simulated and a sample of five diploid individuals was then exported as a Variant Call Format (VCF) file, before being split into 16 chromosomes with sizes identical to those of the *S. cerevisiae* genome. A uniform mutation rate of 1.7e-7 mutations per bp per individual per generation, and a uniform recombination rate of 1e-7, leading to a r = 4.Ne.r = 0.004 were used. The resulting VCF files were used to generate MSMC2's input files using the 'generate_multihetsep.py' script from the 'msmctools' package (version 2018, November 27$^{th}$). Two sets of files were generated: one without callability mask, and one using the callability mask from the Far East Asia population. The MSMC2 program was run on the resulting datasets, first using the default time discretisation scheme (1*2+25*1+1*2+1*3), and then using a scheme with less parameters where the first and last five intervals share a single coalescence rate (1*5+20*1+1*5).

We conducted another set of simulations to assess the impact of background selection on demographic inference. Forward-in-time simulations using the SliM simulator v3.6 (Haller, Galloway, Kelleher, Messer, & Ralph, 2019; Kelleher, Etheridge, & McVean, 2016) were combined with the coalescent simulator ‘msprime’ v1.1.0. As in the neutral case, a population of 10,000 individuals was simulated, using the same genome characteristics and mutation and recombination rates as in the neutral case (L = 12,071,326 bp, u = 1.7e-7, r = 1e-7). In addition, coding regions were defined according to exon annotations from the R64 reference genome (see Genome alignment and gene extraction). Overlapping exons were merged. While recombination was modeled for the full chromosome sequence, only mutations occurring within exons were modelled, with a fitness effect drawn from a negative Gamma distribution with mean equal to -5/N = -5e-4 and a shape equal to 1. All mutations were semi-dominant. After 200,000 generations, the results were saved as succinct tree sequences (Kelleher et al., 2016). The ‘pyslim’ python module v0.700 was then used to ensure that all lineages had a single common ancestor, a process referred to as ‘recapitation’. Five diploid individuals were then randomly selected and the corresponded sub-tree sequence imported into msprime to add neutral mutations. The resulting polymorphic sites were then exported as VCF files. A similar simulation procedure was conducted with a declining demography: after a burn-in phase of 10,000 generations, the population size declined exponentially to 100 individuals at 200,000 generations. The resulting VCF files were processed as in the neutral scenario, split into chromosomes and adding the callability mask of the

Far East Asia population. MSMC2 inference was conducted using a 1*5+20*1+1*5 time discretization scheme.

### Supplementary Results

#### Population phylogeny reflects life history background

To ensure that our populations form well-defined monophyletic lineages, we performed a phylogenetic analysis of the 330 selected *S. cerevisiae* isolates. The resulting tree is consistent with the established scenario of a Far East Asian origin of *S. cerevisiae* (Bendixsen, Gettle, Gilchrist, Zhang, & Stelkens, 2021; Duan et al., 2018; Peter et al., 2018; Wang, Liu, Liti, Wang, & Bai, 2012), with modern non-Asian domesticated yeasts descending from a single shared out-of-Asia event (Peter et al., 2018). Interestingly, the Asian & Domesticated lineages show a distinct ancestry from the Non-Asian & Domesticated lineages, implying that these arose from an independent domestication event, as previously observed by (Fay & Benavides, 2005).

There are a few notable exceptions to the clear distinction between lineages considered Non-Asian & Domesticated and Wild. First, the natural Mediterranean Oak lineage is embedded within the Non-Asian & Domesticated cluster. This is an indication that this lineage is descendent from yeasts historically implicated in industrial processes that have spilled over into natural environments and become feral, a process known to occur frequently (Money, 2018; Wang et al., 2012). Other seeming outliers are the African Palm Wine and African Cocoa lineages which we define as wild, despite being considered domesticated by (Peter et al., 2018). For both these lineages, we argue that it is possible that these do not originate necessarily from the original domesticated stock that gave rise to all the non-Asian domesticated lineages but rather from wild strains that were inadvertently introduced into local ecology through human migration, as is known to underly several other wild populations (Ludlow et al., 2016). Importantly, both African Palm Wine and

African Cocoa fermentation rely on the introduction of *S. cerevisiae* from environmental sources rather than from established domesticated stocks (Djeni et al., 2020; Schwan & Wheals, 2004), and both these lineages appear more related to the wild lineages than to non-Asian domesticated lineages, especially so for the African Palm Wine lineage *(Fig. 1)*. Furthermore, (Ludlow et al., 2016) identified significantly higher levels of genetic variation in African Cocoa lineages compared to the *bona fide* non-Asian domesticated wine lineage, a feature usually associated with populations of a wild rather than domesticated origin. Another striking aspect of the African Cocoa lineage is its paraphyly, revealing three distinct ancestries. This was not noted by (Peter et al., 2018), but such independent origins and high levels of geographic specificity among African Cocoa-associated populations have been reported previously by (Ludlow et al., 2016). A limited interbreeding between local populations is another possible indication for a life history dominated by natural influences, as oftentimes strains are traded and transported more readily as a result of human associations. For these reasons, we shall consider African Cocoa and African Palm Wine as part of the Wild cluster in further analyses.

Combined, our phylogenetic analysis confirms that the populations defined here generally represent clearly distinguishable lineages and, as such, these can be treated as separate populations in population genomics inference methods. Furthermore, the populations can be grouped in three main clusters representing distinct life histories, with one representing a wild origin, the second a domesticated Asian origin, and the third a domesticated and non-Asian origin.

#### Supplementary Tables

**Supplementary Table 1:** List of all selected isolates, their designated population, assembly identifiers, zygosity, ploidy and aneuploidy status (provided as a separate file).

**Supplementary Table 2:** Length of nucleotide alignment for each population after whole genome alignment and after extraction of CDS regions, and the total number of genes recovered from each population.

| Population | Number of Isolates | Original Alignment | Alignment After Filtering | CDS | Number of Genes |
|---|---|---|---|---|---|
| African Beer | 16 | 13,282,799 | 10,191,859 | 7,505,394 | 5,068 |
| African Cocoa | 13 | 12,232,302 | 10,147,887 | 7,480,858 | 5,012 |
| African Palm Wine | 21 | 12,328,332 | 11,046,174 | 8,250,508 | 5,532 |
| Ale Beer | 16 | 12,823,266 | 9,829,821 | 7,179,482 | 4,877 |
| Alpechin | 7 | 11,550,190 | 11,233,073 | 8,338,210 | 5,610 |
| Asian Rice Fermentation | 8 | 12,173,076 | 10,860,914 | 8,067,056 | 5,425 |
| Bioethanol | 32 | 12,463,251 | 10,960,654 | 8,121,915 | 5,462 |
| Dairy | 26 | 12,454,690 | 10,548,655 | 7,843,758 | 5,269 |
| Ecuadorian | 7 | 12,136,055 | 11,347,541 | 8,401,864 | 5,639 |
| Far East Asian | 8 | 12,163,794 | 11,277,937 | 8,356,131 | 5,602 |
| Human French Guiana | 25 | 12,291,185 | 10,752,236 | 8,007,339 | 5,395 |
| Italian Wine 1 | 36 | 12,376,840 | 11,041,381 | 8,245,932 | 5,530 |
| Italian Wine 2 | 35 | 12,344,607 | 11,025,110 | 8,228,014 | 5,524 |
| Malaysian | 6 | 12,121,134 | 11,235,714 | 8,282,951 | 5,570 |
| Mediterranean Oak | 8 | 12,143,626 | 11,260,590 | 8,337,596 | 5,594 |
| Mezcal | 7 | 12,187,688 | 10,096,582 | 7,450,369 | 5,000 |
| North American Oak | 13 | 12,134,523 | 11,370,392 | 8,416,525 | 5,651 |
| Sake | 45 | 12,343,923 | 11,148,616 | 8,294,168 | 5,568 |

**Supplementary Table 3**: Effect of life history, culture method and effective population size (as estimated by $\pi_S$) on the genome average rate of adaptive and non-adaptive non-synonymous substitutions when accounting for population-specific genes.

| | a | | | wa | | | wna | | |
|---|---|---|---|---|---|---|---|---|---|
| Variable | Coefficient | P value | | Coefficient | P value | | Coefficient | P value | |
| (Intercept) | 0.517 | 2.09E-05 | (***) | -17.09 | 0.001 | (***) | 0.083 | 3.91E-05 | (***) |
| Life history, non-Asian & domesticatec | -0.319 | 0.011 | (*) | -14.69 | 0.019 | (*) | 0.056 | 0.010 | (**) |
| Life history, Asian & domesticated | -0.437 | 0.005 | (**) | -23.08 | 0.004 | (**) | 0.076 | 0.005 | (**) |
| Method, repitching | -0.036 | 0.775 | | -4.30 | 0.501 | | 0.006 | 0.787 | |
| Method, starter-based | -0.109 | 0.310 | | -9.58 | 0.094 | (.) | 0.018 | 0.337 | |
| pS | -13.000 | 0.420 | | 149.41 | 0.853 | | 2.151 | 0.440 | |
| Phylogeny (rho) | 2.18E-10 | | | 5.68E-10 | | | 4.22E-10 | | |
| Normality | | 0.780 | | | 0.103 | | | 0.696 | |
| Independence | | 0.216 | | | 0.868 | | | 0.223 | |

**Supplementary Table 4:** Genome average recombination rate estimates, obtained with iSMC using a homogeneous recombination model.

| Population | r | q | r /q |
|---|---|---|---|
| African Palm Wine | 1.55E-04 | 2.25E-03 | 0.069 |
| Alpechin | 4.57E-04 | 4.80E-03 | 0.095 |
| Bioethanol | 8.16E-04 | 3.19E-03 | 0.256 |
| Ecuadorian | 4.00E-04 | 3.60E-03 | 0.111 |
| Far East Asian | 7.94E-04 | 4.77E-03 | 0.166 |
| Italian Wine 1 | 2.00E-04 | 1.31E-03 | 0.152 |
| Italian Wine 2 | 1.90E-04 | 1.24E-03 | 0.153 |
| Malaysian | 3.53E-04 | 4.08E-03 | 0.086 |
| Mediterranean Oak | 1.82E-04 | 2.31E-03 | 0.079 |
| North American Oak | 7.12E-05 | 1.99E-03 | 0.036 |
| Sake | 2.25E-04 | 1.39E-03 | 0.162 |

#### Supplementary Figures

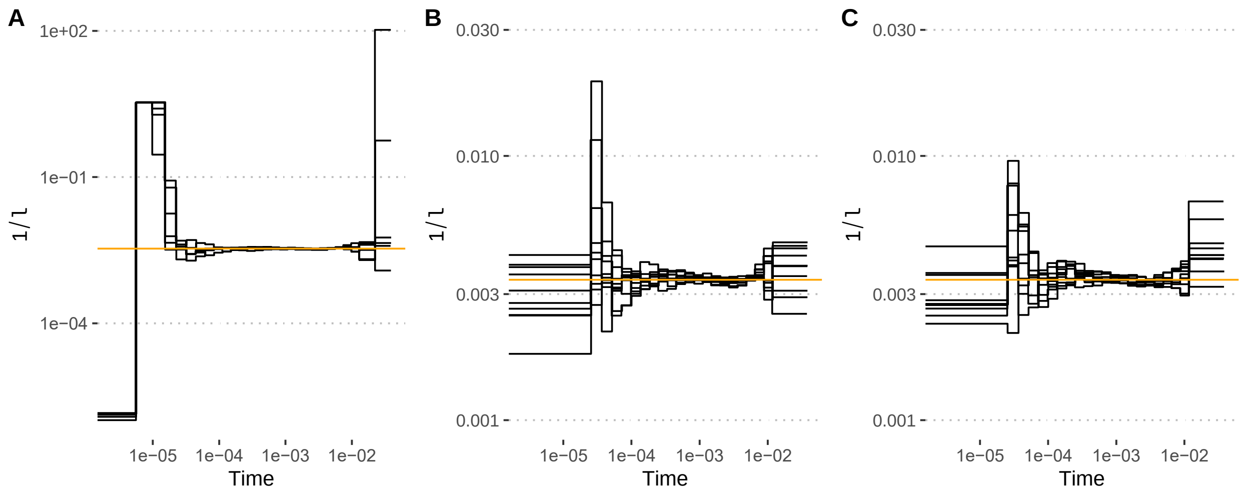


**Supplementary Figure 1**: **Demography recovery from simulated data with yeast genome characteristics, under a neutral scenario and flat demography**. The true demography (constant population size) is plotted in orange. The inferred demography from ten replicates is plotted with black lines. A) Demography inference using MSCM2's default time discretization. B) Demography inference using a 1*5+20*1+1*5 time discretization scheme. C) Demography inference using a 1*5+20*1+1*5 time discretization scheme and callability mask of the real Far East Asia population.

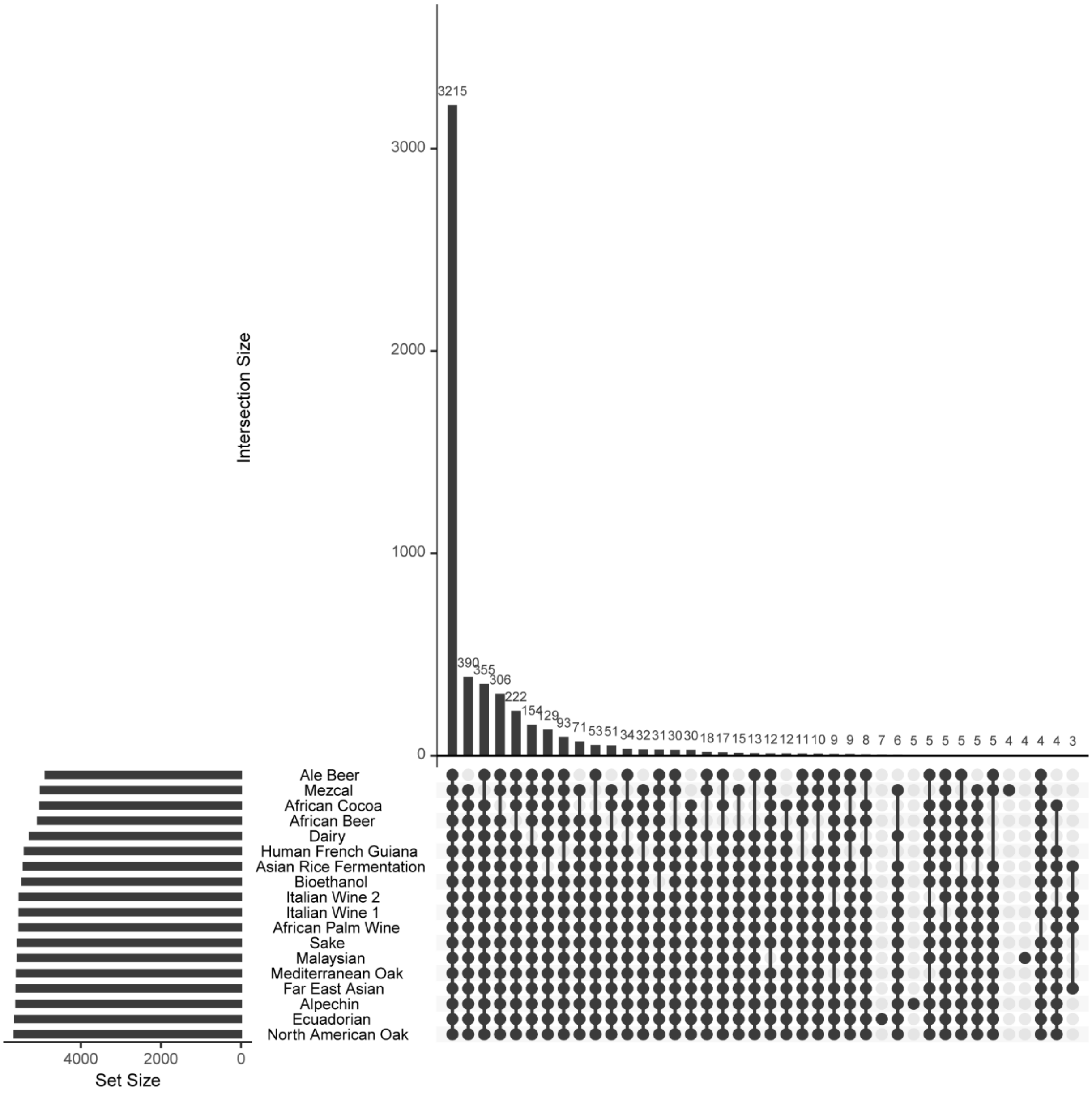


**Supplementary Figure 2: UpSet plot showing the distribution and overlap of genes across different populations.** Diagram summarising the number of genes for all unique intersection set. Corresponding intersection sets are represented with the dots connected by lines in the plot underneath the main bar plot, where filled-in dots signify a population is part of the set, and where a non-included population is shown as a light grey dot with possibly the dark line spanning across it. The set size plotted on the left of the figure represents the total amount of genes recovered from each of the populations. This figure was created using the ‘UpSetR’ package v.1.4.0 (Conway, Lex, & Gehlenborg, 2017).

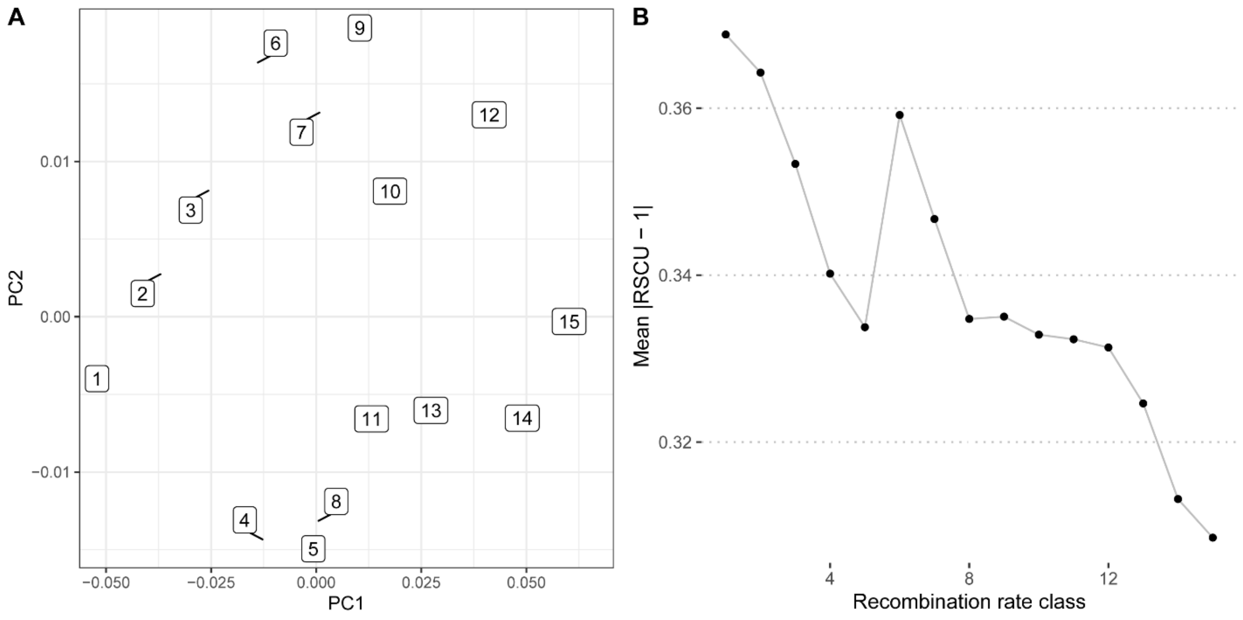


**Supplementary Figure 3: Codon usage analysis with respect to the crossing-over rate.** A) within-group correspondence analysis. Labels correspond to the 15 categories of sites with increasing recombination rate. B) Codon usage bias as a function of recombination rate class. Codon usage bias was measured as the mean relative synonymous codon usage (RSCU) – 1 (absolute value).

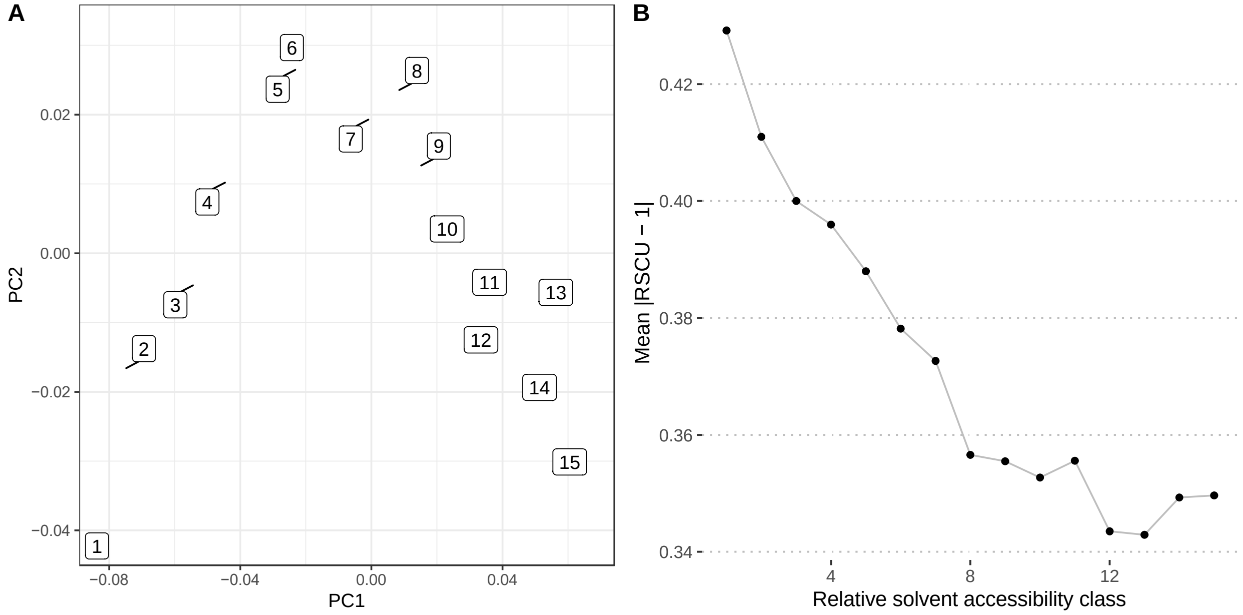


**Supplementary Figure 4: Codon usage analysis with respect to relative solvent accessibility (RSA).** A) within-group correspondence analysis. Labels correspond to the 15 categories of sites with increasing RSA. B) Codon usage bias as a function of RSA class. Codon usage bias was measured as the mean relative synonymous codon usage (RSCU) – 1 (absolute value).
